# LitGene: a transformer-based model that uses contrastive learning to integrate textual information into gene representations

**DOI:** 10.1101/2024.08.07.606674

**Authors:** Ala Jararweh, Oladimeji Macaulay, David Arredondo, Olufunmilola M Oyebamiji, Yue Hu, Luis Tafoya, Yanfu Zhang, Kushal Virupakshappa, Avinash Sahu

## Abstract

Representation learning approaches leverage sequence, expression, and network data, but utilize only a fraction of the rich textual knowledge accumulated in the scientific literature. We present LitGene, an interpretable transformer-based model that refines gene representations by integrating textual information. The model is enhanced through a Contrastive Learning (CL) approach that identifies semantically similar genes sharing a Gene Ontology (GO) term. LitGene demonstrates accuracy across eight benchmark predictions of protein properties and robust zero-shot learning capabilities, enabling the prediction of new potential disease risk genes in obesity, asthma, hypertension, and schizophrenia. LitGene’s SHAP-based interpretability tool illuminates the basis for identified disease-gene associations. An automated statistical framework gauges literature support for AI biomedical predictions, providing validation and improving reliability. LitGene’s integration of textual and genetic information mitigates data biases, enhances biomedical predictions, and promotes ethical AI practices by ensuring transparent, equitable, open, and evidence-based insights. LitGene code is open source and also available for use via a public web interface at litgene.avisahuai.com.

## Main

Genes encode the instructions essential for the synthesis of proteins, which underpin nearly every enzymatic activity and cellular structure required for life [1]. Despite the considerable advances in experimental genetics, our understanding of gene and protein function is far from complete, especially considering the extraordinarily diverse range of biological contexts in which genes and proteins operate [81]. Experimental models capture a tiny fraction of biological contexts, and many genes and proteins remain poorly characterized [43]. This sparse understanding of gene and protein function is exacerbated by human biases. Researchers tend to focus on a small subset of genes, even though whole-genome sequencing has revealed many new genes and their links to diseases [20]. In this manuscript, the term “gene function” will be used to include the function of the DNA-encoded gene itself and its RNA and protein products.

To augment laboratory models, task-specific machine learning models have been developed to predict attributes of genes [58, 63]. These models were trained to perform particular predictive tasks and therefore require extensively labeled data for training. The creation of these labels entails labor-intensive and costly laboratory experiments, which are often not feasible on a scale sufficient for training robust models. This need for large, task-specific training datasets limits the broader applicability and scalability of these predictive models.

In contrast, foundation models are capable of adapting to a wide range of tasks without labeled data [9]. These models, once pre-trained on large unlabeled datasets, can be fine-tuned for various predictive tasks, potentially outperforming task-specific models. A notable advancement in foundation models is the development of Large Language Models (LLMs), which utilize vast bodies of unstructured text. These LLMs enable applications such as few-shot and zero-shot learning [97]. Critically, the knowledge derived from unstructured text in LLMs can be further enhanced by incorporating structured information through techniques like contrastive learning [78, 96] and BootStrap Your Own Latent (BYOL) [24], further improving model performance.

In biomedicine, AI models such as AlphaFold, ESM, BERT, GeneFormer, etc. have led to significant progress in understanding protein-protein interactions, genetic interactions, protein structures, and cellular attributes by leveraging sequence, gene network, transcriptome, or single-cell RNA-Seq data [76, 16, 38, 14]. However, these models exhibit three shortcomings. First, they predominantly harness quantitative data without integrating the rich qualitative insights present in the text of the scientific literature. Second, the ‘black box’ nature of such systems obscures the reasoning behind their output [50], posing significant risks when applied in biomedicine, where decisions can affect patient outcomes. Third, these models have the propensity to amplify human and experimental biases [22], which threatens to perpetuate disparities in healthcare outcomes such as systematic underdiagnosis in marginalized groups.

Here we introduce LitGene, an interpretable large language model designed specifically for gene-related tasks. Beginning with a human-readable text summary for each gene, we utilize an LLM pre-trained on a biomedical corpus to create initial gene embeddings. These embeddings are then refined through contrastive learning, integrating GO annotations to infuse structured biological knowledge into the model. LitGene enables zero-shot learning capabilities and harnesses the wealth of information embedded in the unstructured textual data in the scientific literature related to genes. Importantly, LitGene demonstrates that text-based inputs provide information complementary to sequence and expression data. Our study further demonstrates LitGene’s versatility in predicting gene annotations and associations with disease risk, which showcases its robust zero-shot learning abilities. By supplementing unstructured text with less-biased structured data [16], we also address the knowledge bias inherent in human-led research whereby some genes are extensively studied while others are essentially ignored. LitGene serves to enhance interpretability and reduce bias, significantly improving the utility of AI in gene function prediction in various biological contexts. LitGene is open source and available for use via a public web interface.

## Results

### LitGene Learns Gene Representations From Literature-Based Gene Summaries And GO Terms

We developed LitGene, an interpretable model leveraging the transformer-based BERT (Bidirectional Encoder Representations from Transformers) framework, initially pre-trained on a corpus of PubMed research articles to capture biomedical contexts [26] (Figure 1). To generate gene embeddings, LitGene trains on gene summaries from my-gene.info ([90], Methods) as inputs. However, embeddings generated in this way fail to distinguish genes with different GO annotations (Initial embedding in Figure 1, Supplementary Figure 1). This suggests that the initial LitGene model insufficiently embeds biological contexts, likely due to the similarity in vocabulary and style across summaries.

**Figure 1:**
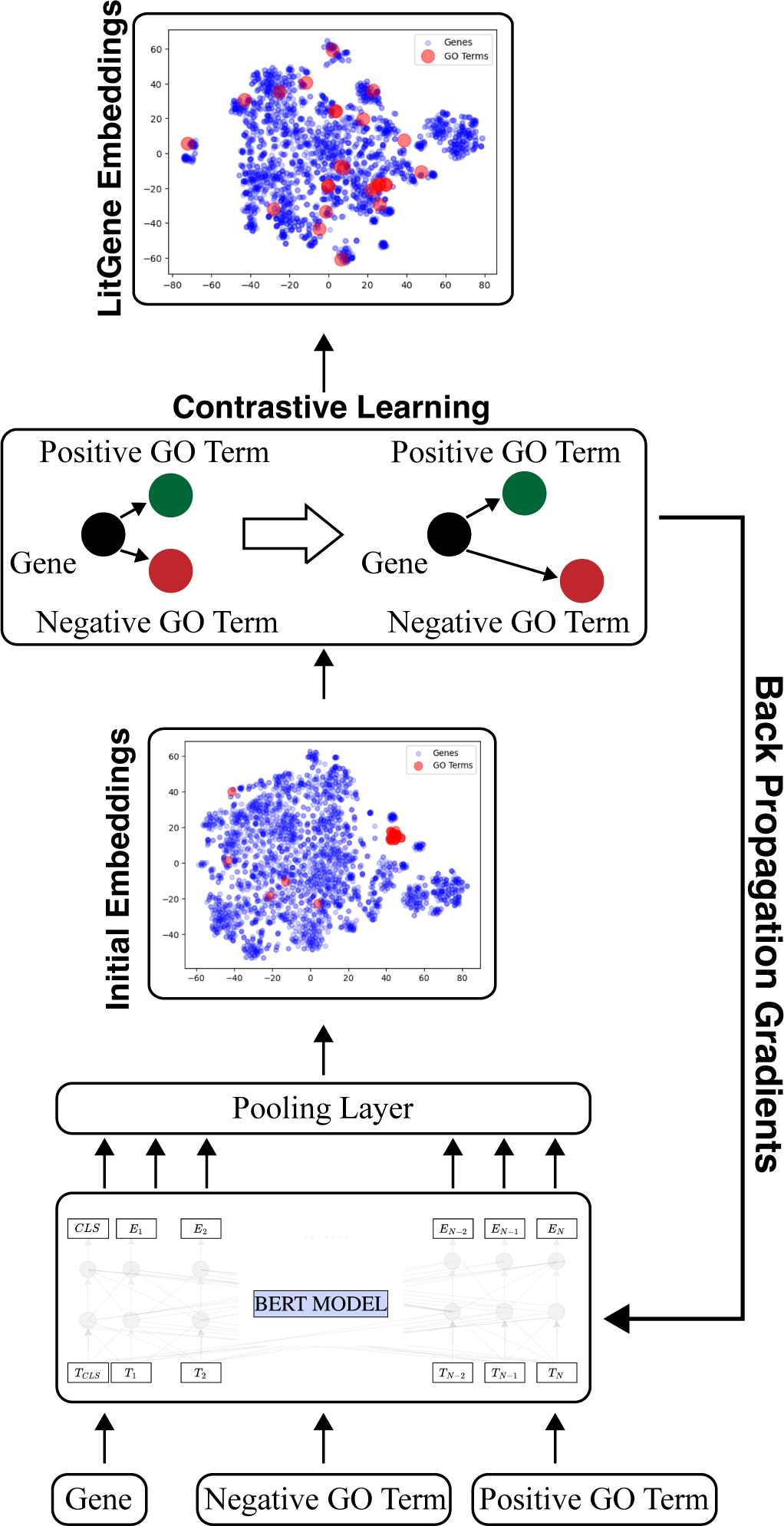
Overview of LitGene Workflow for Gene Embedding Refinement. The initial gene embeddings generated by LitGene-Base are trained on gene summaries, which fail to distinguish genes with different GO annotations (red dots). Many GO terms aggregate together due to similar vocabulary and style in their summaries. Through contrastive learning, these initial embeddings are adjusted by enforcing smaller distances between the gene embeddings and analogous GO terms while increasing the distances from unrelated GO terms, resulting in more accurate and meaningful gene embeddings, as shown in the final LitGene embeddings.

To enhance the ability of LitGene’s embeddings to differentiate genes based on GO annotations, we developed a method based on contrastive learning [12]. This method predicates that embeddings of genes with common GO annotations should converge, whereas those without common GO annotations should diverge. Note that only terminal GO annotations, those at the end of a branch, were used here. Initially, we constructed a knowledge graph of associations between genes and GO terms, comprising 457,000 relationships. We also established filtering criteria to exclude gene/GO terms with duplicate or missing values and to exclude certain words from gene and GO summaries to prevent data leakage (Methods)[86]. This process yielded a knowledge graph linking 21,000 GO terms with 15,000 genes, and a total of 392,000 relationships. We then excluded genes and GO annotations for which we could not obtain summaries, which resulted in a final count of 235,000 relationships.

Subsequently, we co-embedded GO terms and genes using their respective summary descriptions. To encode the knowledge graph of gene-GO relationships into LitGene’s embeddings, we optimized LitGene’s parameters. The optimization minimizes the distance between embeddings of related annotated gene-GO pairs while maximizing the distance for unannotated pairs (Figure 1(b)), which were chosen randomly (Methods). The goal of the optimization is to improve the biological relevance of the generated embeddings. The resultant embeddings can be used to fine-tune downstream gene-, protein-, or cell-related tasks, or for zero-shot predictions.

### Evaluating LitGene’s Predictive Accuracy for Protein Solubility

To assess the effectiveness of LitGene for gene-related tasks, we selected protein solubility prediction as a case study. Proteins perform several critical functions, such as acting as transporters, receptors, pharmacological therapeutics, and enzymes, and many of these functions depend on their solubility [19]. Inferring a protein’s solubility solely from its bioinformatic attributes, such as the number of hydrophobic and hydrophilic residues is often not accurate [44]. Therefore, traditionally the solubility of a protein is determined through laborious experimental methods, including protein expression and purification followed by solubility testing under various conditions, in situ staining, or subcellular localization [84]. This prompted us to investigate whether the text summary associated with a gene could predict the solubility of its protein products.

We benchmarked solubility prediction approaches using data from two studies: one using *in vitro* centrifugation assays that determine whether proteins tend to aggregate or remain soluble [57] and one that experimentally confirms excretion or localization to the cytosol [2] (Methods). These yielded a total of 998 true positive soluble proteins and 966 true positive insoluble proteins. In cases where the two studies disagree on the label of a protein, the centrifugation study was preferred. These benchmarks have been individually assessed by other machine learning approaches [35, 36, 73].

Text-based approaches, including LitGene, demonstrated superior performance compared to methods based on gene expression alone (Figure 2a). LitGene is also benchmarked against amino acid sequence models (see section below). Fine-tuning LitGene specifically for solubility significantly enhanced its performance, achieving an F1-score of 0.84 (Figure 2a). We further benchmarked LitGene against thirteen solubility prediction models specifically trained for this task (Supplementary Table 2). LitGene demonstrated superior performance, exhibiting a 40% increase in F1 score compared to the nearest competitor. This result aligns with trends observed in other domains, where foundation models have matched or exceeded the performance of task-specific models [34, 15].

**Figure 2:**
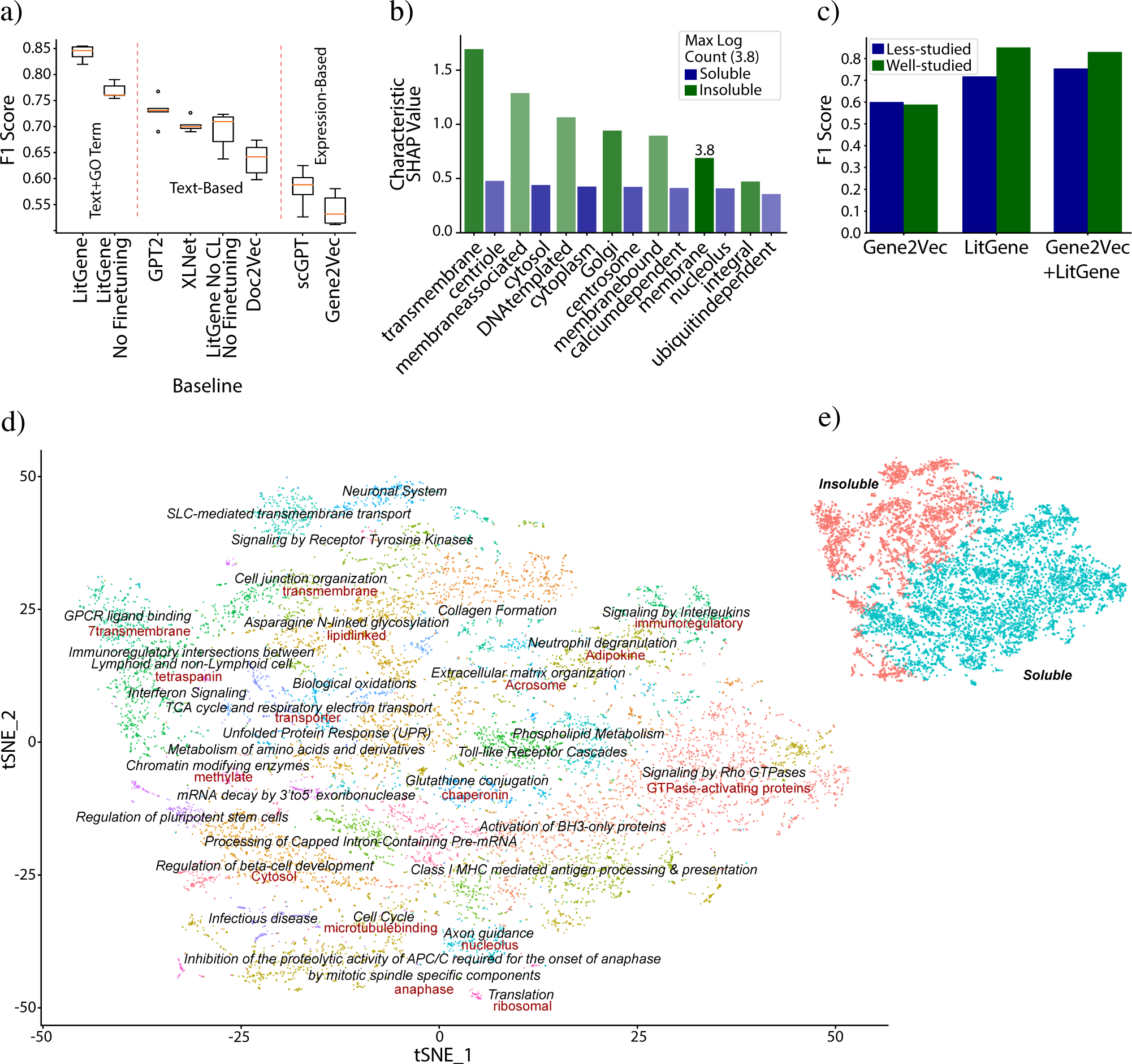
Performance and interpretability of LitGene on solubility benchmark. **a)** LitGene outperform text-based (GPT2, XLNet, Doc2Vec) and expression-based (scGPT, Gene2Vec) models. **b)** Top 7 terms with the highest SHAP values for predicting protein solubility, shown for soluble (blue) and insoluble (green) categories. **c)** Comparative bias analysis of Gene2Vec and LitGene on less-studied and well-studied genes. **d)** A t-SNE plot depicting gene clusters, with pathway enrichment analysis labels and by corresponding important SHAP-derived (frequent) terms in red. An example of interpretative insight: the “GPCR ligand binding” cluster are likely “insoluble” because they are “transmembrane”. **e)** Scatter showing predicted solubility of clusters in (d).

### Interpretability of Model Predictions Using SHAP

To elucidate the decision-making process of LitGene in classifying solubility, we applied SHAP analysis, which quantifies the contribution of each input word to the model’s predictions via an importance score, called the SHAP value (Methods). Despite the binary encoding of classes (insoluble as 0 and soluble as 1), the model identified “transmembrane” as the most important word for the insoluble class (Figure 2b). Transmembrane proteins span the cell membrane, have hydrophobic residues that interact with the lipid bilayer, decreasing their solubility [92]. This finding is notable as the model’s training ensures it has no prior knowledge that “transmembrane” words are directly linked to the *insoluble* class.

We enhanced model interpretability by clustering genes using LitGene embeddings and annotating these clusters with enriched pathways (Figure 2d). Individual clusters were predominantly comprised of members of either the “soluble” or “insoluble” (Figure 2d, 2e). Further, each cluster’s dominant, high SHAP value word—termed the “characteristic word”—provides insight into the model’s classification process (Methods). The word “nucleolus” is associated with the axon guidance cluster, aligning with the expected solubility of nuclear proteins [55] (Figure 2d). Additionally, “7Trans-membrane,” the characteristic word for the GPCR Ligand Binding cluster, describes the attribute of GPCRs traversing the membrane seven times, involving hydrophobic residues and decreased solubility, consistent with most members of this cluster being classified as “insoluble” [92]. Chaperonins (characteristic word) are chaperone proteins that assist in correct protein folding, thus preventing aggregation and promoting solubility [69]. This indicates their role in glutathione conjugation, which maintains protein stability and solubility through redox regulation [54]. GTPases and their regulators, GTPase-activating proteins (characteristic word), are generally soluble because they need to diffuse freely within the cell to interact with their targets [31]. Lastly, “transporter,” the characteristic word for the TCA respiratory electron transport cluster, is consistent with the critical role of these membrane proteins in metabolic processes [11]. Thus, SHAP analysis reveals LitGene’s decision-making process and highlights its potential for prioritizing mechanistic hypotheses for experimental follow-up.

### Evaluation of Potential Data Leakage in LitGene

We rigorously evaluated LitGene’s predictive performance to ensure it was not compromised by data leakage. Data leakage occurs when models inadvertently learn from training data that is irrelevant to the task or not generalizable, leading to overly optimistic performance due to non-representative details, such as keyword frequency. This issue undermines a model’s real-world utility by focusing on data points not fundamentally related to the task [39].

We first examined the potential data leakage by the term “membrane”, frequently appearing in gene summaries (33%). Specifically, we evaluated genes labeled as soluble but with summaries containing “membrane”. Despite this mention, 77% of these genes were accurately predicted as soluble (Supplementary Table 3). This suggests that LitGene assesses the context of “membrane” rather than simply classifying based on its presence. Furthermore, the distribution of SHAP values for “membrane” was centered close to zero (Supplementary Figure 4), suggesting that when “membrane” appeared in contexts unrelated to solubility, it did not significantly influence the model’s predictions. Additionally, we adapted the Cloze test [5], a method originally designed to evaluate language models by predicting words in text blanks, to assess if LitGene could internalize biological concepts such as the interaction between membrane characteristics and protein solubility. Using this test, LitGene accurately completed the sentence “Membrane protein is insoluble in water because of their [MASK1]” with “hydrophobic regions” (Supplementary Figure 5). These tests demonstrate the model’s internalization of relevant biological contexts.

Comparisons with baseline classifiers that depended solely on the keywords “membrane” or “soluble” further validated LitGene’s performance. These classifiers achieved accuracies of 0.75 and 0.52, respectively, which were significantly lower than LitGene’s 0.85 accuracy (Supplementary Table 1) emphasizing its utilization of more complex textual features. Lastly, to generalize this finding, we employed basic feature extraction methods — Count Vectorizer and TF-IDF [88] — to examine if simplistic keyword frequency contributed to class information leakage. Both methods resulted in markedly lower performance compared to LitGene (Table 1, Supplementary Figure 6), supporting the absence of data leakage due to keyword spotting. Collectively, these multifactorial analyses suggest that LitGene’s predictive accuracy in solubility contexts is due to its ability to internalize biological context, not data leakage.

**Table 1:** Comprehensive evaluation of LitGene against multiple baselines (Methods): analyzing 5-fold F1 outcomes for contrastive learning-enhanced gene embeddings across a spectrum of gene prediction tasks (Methods).

| Model | Dosage Sensitivity | BivalentVs Lys4 Methylated | BivalentVs Non Methylated | Tf range | Tf target type | Solubility | Subcellular localization | Conservation (Pearson Corr.) |
| --- | --- | --- | --- | --- | --- | --- | --- | --- |
| Majority Classifier | 0.73 ± — | 0.58 ± — | 0.75 ± — | 0.73 ± — | 0.41 ± — | 0.52 ± — | 0.39 ± — | — ± — |
| GPT2 | 0.73 ± 0.03 | **0.86 ± 0.04** | 0.80 ± 0.11 | 0.67 ± 0.04 | 0.17 ± 0.01 | 0.73 ± 0.024 | 0.77 ± 0.01 | 0.31 ± 0.02 |
| Doc2Vec | 0.73 ± 0.06 | 0.82 ± 0.09 | **0.84 ± 0.05** | 0.57 ± 0.04 | 0.20 ± 0.02 | 0.64 ± 0.03 | 0.69 ± 0.01 | 0.34 ± 0.01 |
| PMC-LLaMA | 0.86 ± 0.05 | 0.77 ± 0.04 | 0.83 ± 0.08 | 0.60 ± 0.04 | 0.07 ± 0.01 | 0.78 ± 0.03 | 0.69 ± 0.01 | **0.55 ± 0.01** |
| XLNet | 0.74 ± 0.06 | 0.84 ± 0.06 | 0.82 ± 0.08 | 0.64 ± 0.06 | 0.12 ± 0.01 | 0.70 ± 0.01 | 0.76 ± 0.01 | 0.40 ± 0.01 |
| Gene2Vec | 0.84 ± 0.03 | 0.84 ± 0.06 | 0.69 ± 0.08 | 0.70 ± 0.06 | 0.19 ± 0.01 | 0.54 ± 0.03 | 0.54 ± 0.02 | 0.50 ± 0.02 |
| CountVectorizer | 0.79 ± 0.03 | 0.84 ± 0.02 | 0.76 ± 0.05 | 0.66 ± 0.05 | 0.07 ± 0.04 | 0.67 ± 0.10 | 0.79 ± 0.01 | 0.05 ± 0.02 |
| TF-IDF | 0.81 ± 0.04 | 0.87 ± 0.04 | 0.61 ± 0.01 | 0.62 ± 0.02 | 0.07 ± 0.03 | 0.67 ± 0.10 | 0.81 ± 0.01 | 0.03 ± 0.02 |
| LitGene - no Finetuning | 0.73 ± 0.08 | 0.83 ± 0.06 | 0.70 ± 0.05 | 0.77 ± 0.04 | 0.07 ± 0.01 | 0.76 ± 0.01 | 0.79 ± 0.01 | 0.48 ± 0.02 |
| LitGene | **0.87 ± 0.06** | **0.86 ± 0.08** | 0.82 ± 0.08 | **0.85 ± 0.04** | **0.48 ± 0.03** | **0.84 ± 0.01** | **0.83 ± 0.01** | 0.53 ± 0.01 |

### Evaluating and Interpreting LitGene on Gene Tasks

We benchmarked LitGene using eight gene-related tasks (Table 1, Supplementary Table 1 for accuracy values). LitGene demonstrated superior performance in five of the eight tasks, particularly Gene Dosage Sensitivity, which is crucial for interpreting the potential effects of copy number variants in genetic diagnostics (Table 1). LitGene also outperformed the baseline predictions (Table 1) in TF Target Type Identification, Protein Localization, and Solubility. These findings suggest textual data are effective for tasks requiring a deep understanding of biological processes and molecular functions (Table 1). However, LitGene performed poorly in Chromatin State and TF Range Predictions, likely due to the limitations of text in capturing certain aspects of gene regulation or the lack of descriptions in existing text for these recently discovered aspects of regulation.

To enhance LitGene interpretability, we analyzed the characteristic words for each of the eight tasks using SHAP (Methods). Figure 3 summarizes the characteristic words for each task. For example, the characteristic word “homeobox” is associated with the bivalent classification, perhaps because homeobox genes are known transcription factor regulators of development, and bivalence allows for a triggered response to developmental cues [18, 32]. The characteristic word “HOXA9” contributes highly to the dosage-sensitive classification potentially because HOXA9 is a homeobox gene whose overexpression is linked to leukemia [95], and dosage changes can cause overexpression.

**Figure 3:**
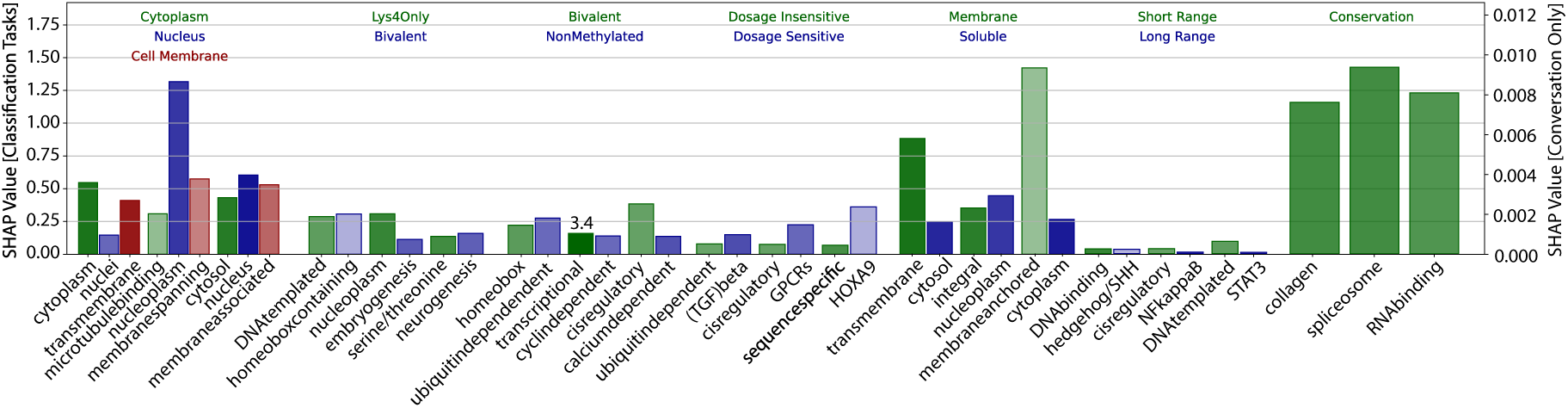
Important words contributing to gene-related tasks. The top three words with the highest characteristic SHAP values (Methods) are shown for each prediction task, indicating their contribution to each output class or value. The alpha of each bar is normalized to the log count of the word’s appearances in the dataset, ranging from a minimum cutoff of log_2_ to a maximum observed count of log_3.4_.

### Contrastive Learning Facilitates Zero-Shot Predictions

We evaluated LitGene’s ability to identify associations between genes and GO terms. We hypothesized that summaries of a gene and associated GO terms would show contextual similarities, which could be quantified by the similarity of their LitGene embeddings. To evaluate this, we used LitGene to embed genes and terminal GO annotations within the same vector space based on their summaries (Methods). In the absence of contrastive learning, the contextual similarity measurements (cosine similarity) from LitGene did not exceed random chance (Figure 4a,b and Supplementary Figure 2; Expanded 4a and 4b in Supplementary Figure 1). However, with contrastive learning, LitGene captures the relationships between genes and GO terms and is able to generalize this capability to novel, unseen GO categories, thus demonstrating zero-shot learning (Figure 4a-c). Further, we evaluated LitGene’s zero-shot prediction ability for KEGG pathway annotations. Remarkably, even without specific training on KEGG pathways, LitGene successfully predicts gene-KEGG pathway associations, further substantiating its zero-shot learning proficiency (Figure 4d).

**Figure 4:**
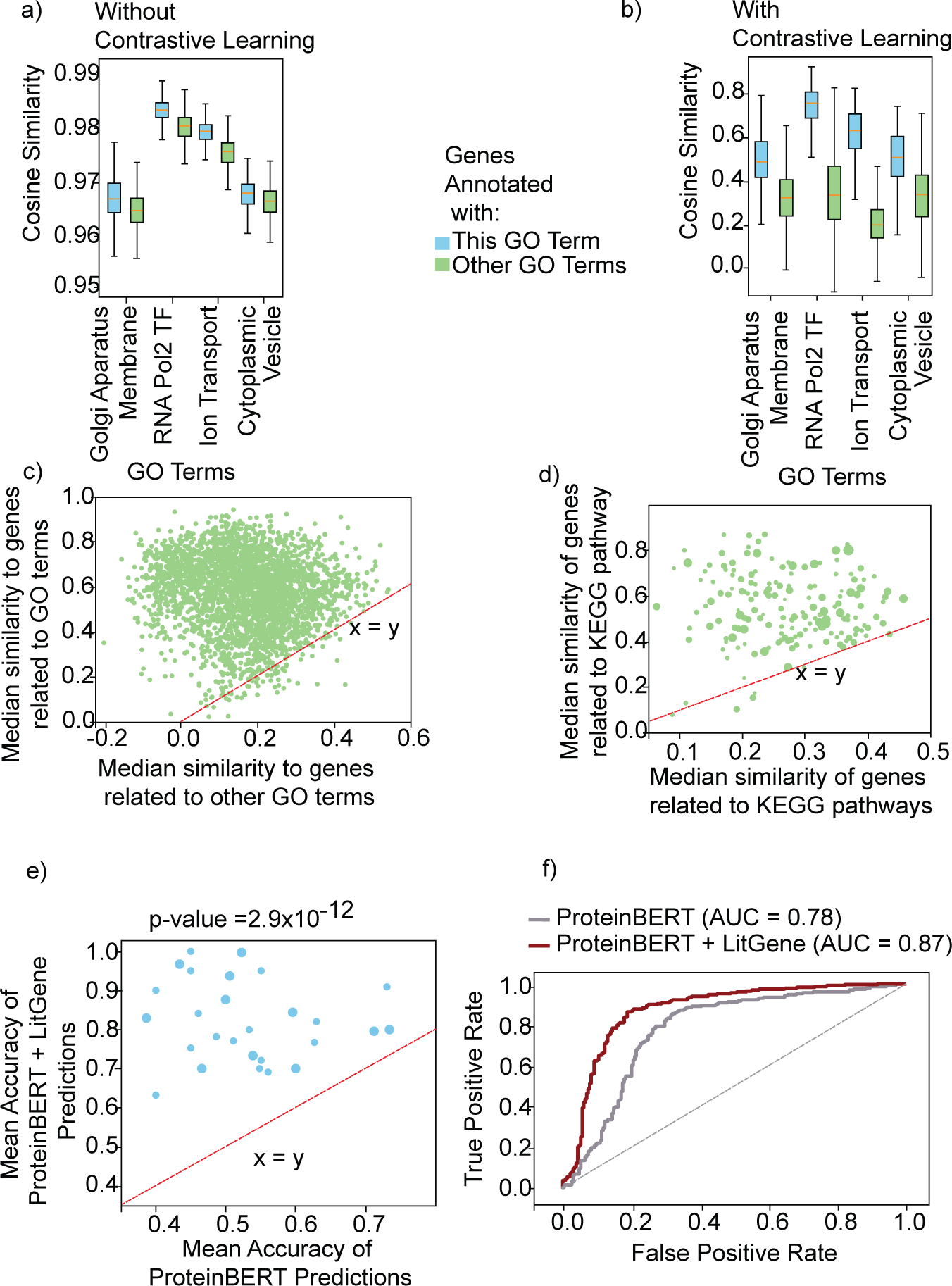
LitGene enables zero-shot learning of novel relationships. **a, b)** Cosine similarity scores between genes and GO terms for a) LitGene-Base (before contrastive learning) and b) LitGene (after contrastive learning). The plots compare scores of genes annotated with the respective GO term to those not annotated. Note the narrower y-axis scale in a. **c,d)** Pathway membership prediction by LitGene trained solely on GO terms. Median cosine similarity of genes related to **c)** GO terms and **d)** KEGG pathways (y-axis) versus genes not related(x-axis). Points above the *x* = *y* line indicate a stronger association with the corresponding GO term or KEGG pathway. **e)** Mean accuracy of 5*−*fold training of KEGG pathways with at least 10 gene relationships. Dots above the *x* = *y* line indicate LitGene+ProteinBERT outperforms ProteinBERT alone. **f)** Performance comparison (AUC) of predicting KEGG pathways comparing LitGene+ProteinBERT versus ProteinBERT alone.

We hypothesized that combining LitGene’s text embeddings with embeddings derived from amino-acid sequences would enhance pathway prediction. We developed a supervised multimodal predictor that merges LitGene embeddings with those from ProteinBERT [10], a model trained on amino acid sequences (Methods). This multimodal approach indeed surpassed the performance of predictors using ProteinBERT embeddings alone (Figure 4 e, f), indicating that textual information meaningfully complements data derived from amino acid sequences.

Although zero-shot learning failed for some GO and KEGG terms due to limited knowledge (Discussion), LitGene’s demonstrated ability opens new possibilities in biology and medicine, especially in contexts lacking labeled data, such as personalized medicine (n=1).

### Zero-Shot Prediction of Disease Risk Genes

We asked if LitGene could identify risk genes of a given disease. Identifying the genetic bases of rare, polygenic, and complex diseases is challenging because rare diseases often lack sufficient cases for effective genome studies [75]. Complex or polygenic diseases, such as schizophrenia, obesity, and hypertension involve intricate interactions between multiple genes and environmental influences [87].

We hypothesized that LitGene’s zero-shot capability could infer disease risk genes by leveraging contextual similarity between disease and gene function descriptions. We collected disease-gene relationships from three datasets, namely DisGeNET [62], MGI [45], and PsyGeNET [30]. Without training LitGene specifically on disease datasets, we calculated the cosine similarity of LitGene’s embeddings of disease and gene descriptions. LitGene demonstrated zero-shot prediction by showing similarity between disease and gene embeddings, in all three datasets, suggesting a generalizable approach to infer disease risk genes (Figure 5a and Supplementary Figure 7).

**Figure 5:**
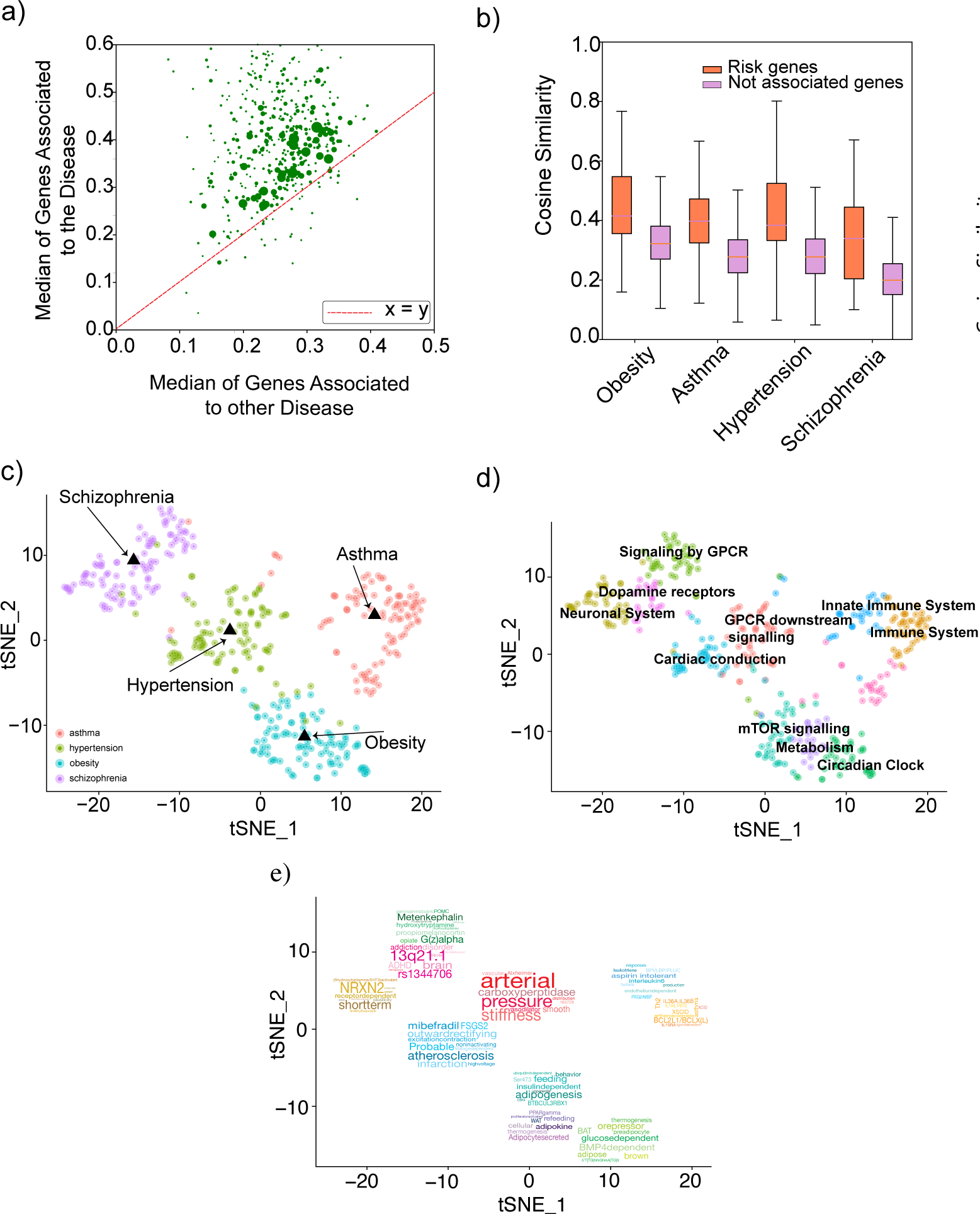
Zero-Shot Prediction and Disease Association Analysis Using LitGene. **a)** Median cosine similarity of genes associated with a specific disease (y-axis) compared to those associated with other diseases (x-axis). Points above the *x* = *y* line indicate a stronger association with the specific disease. **b)** Cosine similarity between gene embeddings and disease-specific embeddings, comparing genes annotated with each disease (red) versus those not associated (purple). **c-e)** t-SNE plots in the same embedding space: **c)** Top novel 100 predicted genes by LitGene not annotated in gene-disease databases, colored by disease label: asthma (red), hypertension (green), obesity (blue), and schizophrenia (purple). **d)** Enriched biological pathways within each disease-specific gene cluster. **e)** Clusters with the most important words contributing to their prediction according to SHAP analysis.

Next, we explored the potential of this predictor to find new gene-disease associations, focusing on asthma, hypertension, obesity, and schizophrenia. We compiled gene sets known to be associated with these diseases, referred to as their respective “ground truth genes”. The embeddings of obesity, asthma, hypertension, and schizophrenia were significantly more similar to their respective disease ground truth genes than to random genes, suggesting LitGene could discriminate risk genes from random genes (Figure 5b). For each disease, we next identified genes that exhibited a higher or similar level of cosine similarity to the respective ground truth genes (Supplementary Table 4 in Excel file). We clustered these genes, which were not previously reported to be linked with their respective diseases, and identified enriched pathways and characteristic words using SHAP (Figures 5 c-e; Methods). This analysis yielded clues regarding the nature of the potential associations between the genes and disease, as shown in Table 2.

**Table 2:** Summary of characteristic words, with their respective diseases, pathways, and descriptions.

| Characteristic Word | Disease | Pathway | Description |
| --- | --- | --- | --- |
| NRXN2 | Schizophrenia | Neuronal System | NRXN2 is crucial for forming synapses and balancing brain signals. Disruptions can impair brain communication and stability. [23] |
| Adipogenesis | Obesity | mTOR signaling | Adipogenesis is relevant to obesity, promoting cell growth and lipid metabolism. [41] |
| BMP4dependent | Obesity | Circadian clock | BMP4 is involved in adipose tissue differentiation and energy balance. [29] |
| Interleukin6 (IL-6) | Asthma | Innate Immune System | Elevated IL-6 cytokine levels contribute to chronic inflammation and airway remodeling. [68] |
| Th2 | Asthma | Immune System | T-helper type 2 cells play a role in allergic inflammatory responses characteristic of asthma. [21] |
| Arterial, pressure, stiffness | Hypertension | GPCR Downstream Signaling | These words link blood pressure regulation to vascular function. Arterial stiffness is a predictor of heart failure and hypertension. [44] |
| Mibefradil | Hypertension | Cardiac Conduction | T-type calcium channel blockers are used to manage hypertension by reducing cardiac workload and improving arterial compliance. [80] |

### Anchoring Gene-Disease Predictions in Published Research

We tested LitGene’s ability to identify publications with abstracts contextually similar to descriptions of genes and diseases. We embedded abstracts from 108,122 PubMed articles using LitGene and jointly clustered the embedded publication abstracts, gene descriptions, and disease descriptions. The resulting clusters were then plotted separately for visualization, with numbers assigned to each cluster, allowing one to match clusters across plots (Figure 6a,b and Supplementary Figure 3). The co-clustering revealed articles and gene groups that were contextually similar (occurred within the same cluster). For instance, a set of genes enriched in sperm motility, fertilization, and reproduction was clustered with articles about reproduction (Figure 6c, d). Similarly, a set of genes related to immune pathways clustered with articles on immunity, coronavirus, and inflammation (Figure 6e, f). Such relationships were also observed between other genes and articles (Supplementary Figure 8). Therefore, although LitGene was trained for gene descriptions, it could also embed Pubmed articles and uncover themes shared with gene descriptions.

**Figure 6:**
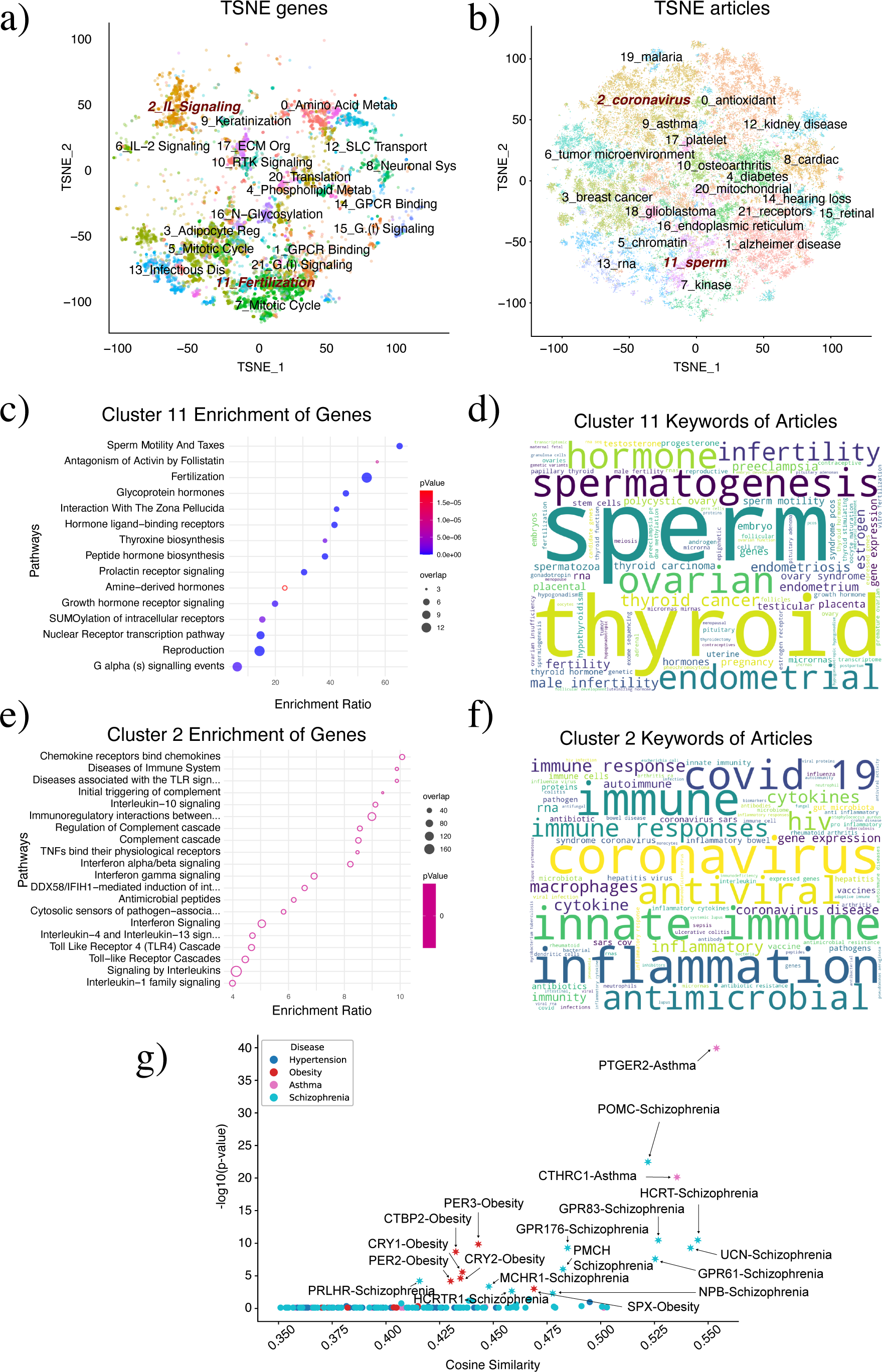
Automated method to gauge literature support for LitGene predictions. **a,b)** TSNE plots of gene (a) and article (b) embeddings. Each colored dot represents a gene or an article, with clusters formed unsupervised. In (a), labels indicate pathways enriched in each gene cluster. In (b), labels represent keywords/topics associated with each article cluster. The labels demonstrate the thematic grouping of gene or article clusters based on their annotations or content. Cluster numbers preceding the labels indicate matched clusters for genes and articles, highlighting the relationship between gene functions and article themes. **c)** Enrichment ratio (observed/expected overlap) and p-value (color gradient) of enriched pathways of genes in Cluster 11. **d)** Word cloud of top topics from articles in Cluster 11. **e)** Enrichment ratio and p-value of pathways for genes in Cluster 2. **f)** Word cloud of top topics from articles in Cluster 2. **g)** Cosine similarity (x-axis) between the gene and the disease embeddings and significance (p-value) of overlap of publications related to genes and diseases (y-axis), with pairs significant after Benjamini-Hochberg (BH) multiple-hypothesis correction marked by asterisks.

Next, we evaluated whether gene-disease relationships inferred by LitGene are corroborated by existing studies. We defined corroboration as a significant overlap in publications related to the gene and the disease. To test this, we used contextual (cosine) similarity to identify publications related to each gene, and separately to each disease. We then calculated the significance of the overlap using the right-tailed hypergeometric test (Methods). This analysis identified 19 gene-disease pairs inferred by LitGene that are significant after Benjamini-Hochberg (BH) multiple-hypothesis correction and are also supported by the literature (Figure 6g).

For example, the candidate gene most highly ranked by LitGene to be associated with asthma was PTGER2. Although neither the Mouse Genome Informatics Database (MGI) [45] and PsyGeNET [30] explicitly annotate PTGER2 and asthma as associated, this relationship is supported by genetic, expression, and functional studies in the literature [42, 53, 70, 13]. Polymorphisms in PTGER2 are linked to asthma susceptibility [42, 53]. PTGER2 is a receptor for Prostaglandin E2 (PGE2), which has multiple effects on immune responses in asthma [70]. Additionally, PTGER2 has been identified as a potential molecular biomarker for asthma [13]. A second example of a gene-disease relationship predicted by LitGene is between hypocretin (HCRT) and schizophrenia. Again, this relationship is strongly supported by literature. Schizophrenia patients exhibit significantly higher plasma hypocretin levels and epigenetic alterations in the system [51, 28]. Hypocretin plays a crucial role in regulating sleep-wake cycles, attention, cognition, and energy balance, all of which are often disrupted in schizophrenia. These examples indicate LitGene’s zero-shot ability to infer gene-disease relationships supported by literature.

Using this approach, LitGene can generate inferred gene-disease relationships in an automated manner, augmenting and potentially enhancing manual literature searches. Further, this method could be used to restrict inferences made by LitGene to those with support from the scientific literature. Beyond gene-disease relationships, LitGene has been extended to infer relationships between other biological entities, such as disease-drug and gene-drug interactions. These tools are accessible through the LitGene web interface.

### Mitigating Bias in Gene Representation with Multi-Modal Learning

Human understanding of genes is biased, with some genes researched extensively due to their associations with disease or certain functions amenable to experiment, while many remain understudied. This disparity is often compounded by biases in genetic research and publication biases [74]. AI models trained on such data are at risk of perpetuating or even amplifying these biases [72]. Quantifying human biases is challenging [82], we, therefore, focus on addressing the potential bias stemming from limited knowledge. We demonstrated the impact of this bias on AI through two analyses.

In the first analysis, we used the summary lengths of KEGG pathways as proxies for their knowledge levels, to approximate the accuracy of LitGene on well-studied versus less-studied KEGG pathways. As expected, LitGene’s ability to ascertain contextual similarity via zero-shot learning improved with the availability of more information, suggesting that a similar bias would occur in predictions of well-studied vs lesser-studied pathways.

In the second analysis, we evaluated model performance for solubility predictions among genes with limited online information. The results demonstrated poorer performance for these genes (Figure 2c), reinforcing the presence of knowledge bias. We hypothesized that a multi-modal approach that combines unstructured data with structured data could mitigate the impact of knowledge bias. By integrating LitGene’s knowledge embeddings with structured Gene2Vec embeddings, which are derived from gene expression data that is less likely to be affected by human knowledge biases, we narrowed the performance gap for under-researched genes (Figure 2c). This provides evidence that incorporating representations from non-text modalities can help offset biases inherent in text-based data. Furthermore, the complementary nature of text and expression data, along with our demonstration of complementarity between text and protein sequence data (Figure 2c, 4f), supports the broader potential of integrating text and non-text modalities for improving performance and reducing bias.

### A Public Web Interface for LitGene

We provide an accessible web interface for biologists and clinicians who wish to use LitGene. Users can input any gene, disease, or pathway, and the interface outputs a predicted list of top-related genes, diseases, and pathways. Additionally, users can submit any text summary, including new functions, and the website will output any gene, disease, or pathway predicted by LitGene as related to the summary. Furthermore, users can request the interface to identify published articles corroborating LitGene’s predictions, thereby increasing the reliability of the AI predictions. See Supplementary Figures 9, 10, and 11 for example website outputs.

## Discussion

This study aims to develop a model that integrates unstructured textual data with structured biological data to improve gene-related predictions and interpretations. To that end, we introduce LitGene, a foundation model for gene embedding. This goes beyond current biomedical AI models that predominantly utilize quantitative data but do not leverage insights in scientific literature. Previous studies have incorporated GO annotations with protein-sequence-based embeddings [96], whereas our approach utilizes text as the input and contrastive learning to internalize GO-gene network information to refine text embeddings. LitGene enhances the interpretability of outputs, addressing the ‘black box’ nature of many AI systems in biomedicine. This transparency in the decision-making process increases the reliability and applicability of AI in critical biomedical and clinical decisions. LitGene demonstrates the utility of the foundation model by its zero-shot ability and illustrates that integrating structured and unstructured data mitigates knowledge biases prevalent in the literature. Our results are consistent with observations in other domains where the capabilities of foundation models meet or exceed the performance of task-specific models and pave the way for more equitable and comprehensible application of AI to biomedical problems. LitGene is an open-source project that can be modified, fine-tuned, and extended by researchers. By analyzing genomic and textual data, LitGene holds promise for generating and evaluating hypotheses about gene functions, such as oncogenic potential, synthetic lethal interactions, and genetic mechanisms of diseases. It can also generate hypotheses regarding treatment responses, and disease associations, drug-gene interactions, potentially accelerating drug discovery by repurposing existing drugs and identifying disease mechanisms.

We also demonstrated an automated method to gauge literature support for LitGene predictions, which could be used to enhance the model’s reliability. However, this approach may limit the model’s ability to infer relationships between entities that are not yet well-studied in the literature. Therefore, we recommend using LitGene both with and without literature-based restrictions. Retrieval-Augmented Generation (RAG) [49] is another approach that integrates external documents – and could similarly integrate published articles [91] like LitGene’s literature grounding approach – with embedding techniques to find contextual similarities, although RAG is primarily used for natural language understanding and generation.

This proof-of-concept study has several limitations. First, the lack of standard benchmarks in gene-related tasks (analogous to the GLUE benchmark in natural language processing [83]) hampers our ability to compare our results against a universal metric, potentially leading to the over- or under-estimation of model performance. To mitigate this, we evaluated LitGene across a diverse set of tasks, some of which have been employed in other foundational models [76, 85]. The consistently enhanced performance of LitGene across various tasks supports the benefit of adding textual data to current AI models. Second, the quality of our zero-shot predictions is dependent on the quality of the pathway and disease summaries used. These summaries may be incomplete, noisy, or may not effectively address or describe certain aspects of biological function, which may compromise the model’s performance in certain applications, as we observed for Chromatin State and TF Range Predictions. A multimodal approach is by nature well-suited to addressing weaknesses in any individual modality. We enriched our model’s embeddings using gene annotations and employed contrastive learning techniques, which are known for their robustness against incomplete or misleading information [40]. Additionally, LitGene’s reliance on pre-trained language models introduces the risk of information leakage. To mitigate these risks, we conducted analyses to evaluate the impact of information leakage. For instance, in the solubility task, we demonstrated that the model’s accuracy was not solely due to word association biases. Additionally, LitGene embeds genes based on summaries rather than names, reducing the likelihood that specific text used during pre-training unduly influences gene-specific predictions.

There are also ample opportunities for improvement and additional use cases, including predicting risk genes for a given disease, gene targets of a given drug, drugs that might be effective for a given disease, or the side effects of a given drug. Future versions of LitGene could take advantage of the rich information in GO hierarchies to refine gene embeddings. Further, significant improvements were often achieved through fine-tuning. This suggests the potential for developing a unified approach that integrates various tasks into a single prompt-based system, optimizing the algorithm across all tasks concurrently. LitGene’s performance may be enhanced further by entirely different approaches. For example, complex network structures may be more effectively processed using Graph Neural Networks (GNNs). Finally, LitGene is amenable to a variety of downstream tasks in addition to the use cases demonstrated here. LitGene embeddings could be used to predict protein-protein interactions (PPIs), identify synthetic lethal interactions, find targeted therapies for genetic disorders, and formulate new precision strategies for cancer treatment. LitGene represents a step forward in integrating textual and biological data, increasing the reliability of AI by anchoring it on scientific literature, and providing a foundation for future innovations in biomedical AI.

## Methods

### LitGene Backbone - Background

Feature-based and fine-tuned pre-trained language models have shown state-of-the-art performance for various NLP tasks [61, 64]. Fine-tuning models such as BERT seek to minimize the number of task-specific parameters using pre-trained parameters on labeled data [15]. The BERT training consists of two stages: pre-training and fine-tuning. The model is first pre-trained using two self-supervised tasks: masked language modeling (MLM, where the words are masked and the model predicts those masks) and Next Sentence Prediction (NSP, where the model attempts to distinguish whether a given pair of sentences are adjacent or not). BERT variations have also been applied (either by pre-training or fine-tuning) to various biomedical text datasets [27, 47, 94, 3, 60, 4]. LitGene seeks to utilize the abundant human knowledge learned by these models to predict cellular and genetic properties. We designed LitGene to be adaptable to a wide range of BERT-based models (see Fine-tuning section below).

### Gene Summary Pre-processing

The gene summary dataset was curated from two publicly accessible databases. We obtained the gene descriptions from the mygene.info database [90] and the gene function summaries were downloaded from UniProt. Through a manual inspection, we found that some gene summaries contained the underlying values of certain keyword labels such as *pubmed id*, *author’s name*, and *isoform id* that were identical among genes with similar functions. The model captured this information and thus found shortcuts to make predictions based on unwanted content. Consequently, we eliminated such identifying keywords that introduce bias using regular expressions. After preprocessing, we found that some of the less-studied gene summaries have fewer words that do not specifically discuss the genes. These summaries are also found to be the same for all less-studied genes. To ensure the predictions are based on the content of the summary and not on other biologically irrelevant text, we further removed genes described by fewer than 10 words and genes having summaries identical to the summaries of other genes.

### Gene Level Embeddings

We derive gene embeddings by supplying text summaries of the desired genes to LitGene where the text will be tokenized via the selected backbone encoder’s tokenizer. After tokenization, each gene summary is represented as a list of *N* tokens. As a preliminary step, the tokens are passed to the pre-trained BERT model to generate contextualized token representations (i.e. embeddings) where an embedding vector, *E_i_* embeds each token, *T_i_*. Each sentence *S* is represented with *N* contextualized token representations, where *∀i ∈ {CLS,* 1, 2*, …, N }, E_i_ ∈* R^768^. We aim to synthesize an embedding *E_g_*, that transforms *E* into a feature space R^768^ (Figure 1). To accomplish this, we use the pooling strategies introduced in [67] to combine contextualized representations into a fixed-length representation. Three different strategies were mentioned: CLS pooling, mean pooling, and max pooling. For CLS pooling, only the embedding of the classification token *E_CLS_* is considered. Mean pooling is computed by taking the mean of all contextualized representations. Finally, Max pooling is the element-wise maximum of all token representations. The feature vector obtained after pooling has a dimensionality of R^768^.

### Encapsulating GO-Gene Associations via Contrastive Learning

Contrastive learning is a self-supervised machine learning approach that utilizes the similarities between points to instruct model training. Here, we leverage CL to find a space that combines the representations of semantically similar genes closer together, while pushing apart dissimilar genes. Semantically similar genes are identified as the genes that share a GO term. Our approach involves co-embedding genes and GO terms into a shared embedding space. We utilize 457,000 relationships, however, removing entries with missing values and considering relevant relationships that strictly include the genes in our datasets reduces the final count to 296,000 confident relationships. The remaining relationships, *R*, comprise a total of 21,000 GO terms and 15,000 genes that are intended to be co-embedded into the shared space. Of these relationships, 61,000 gene-GO relationships were hidden to evaluate the model training comprising 3000 unseen GO terms. This approach embeds GO terms explicitly and could predict GO-gene relationships for unseen GO terms and hence facilitate zero-shot learning of the unseen GO terms.

Formally, the CL approach aims to bring anchor genes (*g_a_*) closer to their corresponding GO terms (*t_p_*, the positives) in the shared embedding space *E*, while maximizing the distance between *g_a_* and unrelated GO terms (*t_n_*, the negatives). The loss function is defined as:

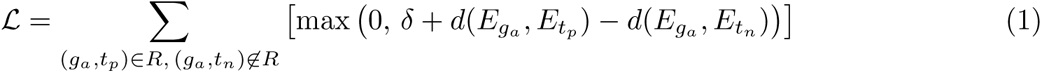

The margin *δ* (0 *< δ <* 1) ensures that the positive pairs are closer to the anchor than the negative pairs.

### LitGene Fine-Tuning

Unlike other gene representation methods where the embeddings are one-size-fits-all, LitGene learns gene embeddings that are customized for each task. The customization is done through fine-tuning LitGene parameters using the labeled data for the desired tasks. The labels can range across different types such as categorical classes (protein localization), gene conservation values, and Gene Interactions or GO annotations. LitGene can also utilize external gene knowledge learned via other gene representation models such as Gene2Vec [17]. The external knowledge is integrated via late fusion during fine-tuning to generate composite embeddings. To fine-tune LitGene, we added a task-specific additional output module on top of the final encoder layer. The model is fine-tuned by minimizing a specific loss function that is determined based on the task. LitGene minimizes the Mean Squared Error (MSE) in regression tasks, Binary Cross-Entropy (BCE) for binary classification tasks, and Cross-Entropy for multi-class classification tasks.

### Zero-Shot Learning of Pathway Annotations

To create Figure 4, we validate the model’s ability to produce zero-shot predictions of pathway annotations (GO, KEGG, etc.) without further training. We produce zero-shot predictions from LitGene by inferencing the model for embeddings of unseen pathways where the pathway embeddings are then compared to the gene embeddings to find analogous gene-pathway relationships. Formally, let 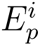 and 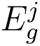 respectively be the embedding of the pathway *i* and gene *j* where both embeddings are obtained from text summaries fed to LitGene. We refer to the *i*-th pathway and *j*-th gene to be analogous if the cosine similarity between their embeddings is higher than a threshold, 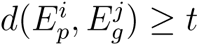.

### Gene-Disease Association

We extended our analysis to explore gene-disease associations using the Mouse Genome Informatics Database (MGI) [45] and PsyGeNET [30]. MGI offers a comprehensive dataset linking 21,671 genes to 30,170 diseases, traits, and phenotypes in both humans and mice. PsyGeNET is a specialized resource for psychiatric disease research, providing associations between 1,549 genes and 117 psychiatric disease concepts, covering disorders such as depression, bipolar disorder, schizophrenia, and substance use disorders. We first extracted textual summaries of genes and diseases from UniProt and Wikipedia. We generated embeddings for these summaries using LitGene. We calculate the cosine similarities between gene and disease embeddings to measure the closeness of their embeddings. The cosine similarity scores served as a quantitative metric of association strength between genes and diseases. High similarity scores indicate a strong association, suggesting that the gene and disease share common biological contexts and functionalities as captured by their textual summaries.

### Gene-Disease Association Profiling Using LitGene and Pathway Enrichment

To uncover patterns within gene-disease associations, we focused our analysis on 400 genes that represented the top 100 genes with the highest cosine similarity scores from LitGene embeddings but that had not been previously associated with each of four diseases in those databases: Asthma, Hypertension, Obesity, and Schizophrenia. This resulted in a cohort of 400 genes.

To create the plot in Figure 5c, we used an approach similar to that used for single-cell data analysis in Seurat [33]. Specifically, Principal Component Analysis (PCA) was performed on LitGene embeddings, reducing the dataset to its top 50 principal components. A k-nearest neighbor (KNN) graph was constructed based on the principal components. This was converted into a shared nearest neighbor (SNN) graph to genes with shared neighbors. Clusters were identified using the Louvain algorithm on the SNN graph with a resolution parameter of 0.5. The resulting clusters were visualized using t-SNE. The clusters were distinctly color-coded by their respective disease labels as shown in (Figure 5c). To create the plot in Figure 5d, we also employed WebGestaltR for a pathway enrichment analysis of each gene cluster. This provided us with a biological interpretation of the clusters, mapping genes to overrepresented pathways.

### Anchoring Gene-Disease Predictions in Published Research

To create Figure 6, we co-clustered genes, diseases, and articles using their LitGene embeddings. Gene and disease summaries were used to obtain LitGene embeddings while PubMed Central (PMC) article abstracts were utilized for article embeddings. Articles were sourced from PMC by identifying publications that mentioned genes, diseases, or drug names from our own database within their abstracts. This initial search yielded over 2 million articles. To refine this list, articles were then filtered based on citation counts, with only those cited more than three times being selected. This process resulted in a final dataset comprising 108,122 articles.

For co-clustering, we combined the LitGene embeddings of genes, diseases, and articles into a single integrated matrix. We then identified clusters and visualized them using t-SNE, following the same procedure described for clustering gene embeddings alone (see sections above). Specifically, we performed PCA on the integrated embedding matrix, constructed a KNN graph, converted it into an SNN graph, and identified clusters using the Louvain algorithm. The resulting clusters were visualized using t-SNE. Although genes, diseases, and articles were clustered jointly, they are plotted separately for ease of display. Each instance of the combined embedding is plotted in a separate scatter plot: genes (Figure 6a), articles (Figure 6b), and disease (Supplementary Figure 3). The cluster numbers remain consistent and matched across gene, disease, and article clusters.

For a given cluster, enriched pathways were identified using the gene sets within the cluster via WebGestaltR’s pathway enrichment analysis. Figure 6c,e display enriched pathways for clusters 2 and 11. In Figure 6a, clusters are labeled based on the most significantly enriched pathway in each cluster.

Top topics within the articles of each cluster were identified using KeyBERT [25]. KeyBERT was applied to extract important topics from the articles, with the input being the text of each article’s abstract within the cluster (Figure 6d,f). The clusters are labeled based on the topmost topics in each cluster.

Finally, we assessed whether LitGene-inferred disease-gene pairs could be corroborated by published articles. For this literature-based validation, we tested the hypothesis that articles related to a specific gene significantly overlap with articles related to a specific disease. Specifically, we identified the top 1,000 articles with the highest cosine similarity scores to each gene and each disease. Cosine similarity was computed based on the LitGene embeddings of the articles, genes, and diseases.

The significance of the overlap between the sets of articles was then evaluated using a right-tailed hypergeometric test. To control for multiple hypothesis testing across the numerous gene-disease pairs, we applied the Benjamini-Hochberg (BH) correction [6] to adjust the p-values, with a significance threshold set at 0.05 after correction(Figure 6g). In interpreting the results, a significant overlap suggests that the inferred disease-gene pair has substantial literature support.

### Mitigating Bias Using Multimodal Approach

To create the plot in Figure 2c, we integrate the LitGene and Gene2Vec embeddings, which are based on gene expression patterns. This integration was achieved by fusing vector embeddings from both models. The resulting integrated embeddings were then used for prediction, aiming to reduce knowledge bias inherent in gene expression data. In this context, gene expression data does not exhibit the same bias as knowledge-based data, suggesting a compensating effect for the ascertainment bias present in text data. We evaluated the extent of bias mitigation by assessing the model’s performance on solubility predictions.

### Integrating Textual Knowledge with Protein Sequences

To create the plot in Figure 4e,f, we applied a multimodal learning approach combining text and sequence features of proteins/genes. This analysis involved using LitGene to incorporate textual knowledge with sequence features extracted by ProteinBERT [10]. Sequence features were derived using the ProteinBERT model, utilizing the amino acid sequence of the dominant protein for each gene. ProteinBERT embeddings (originally *E ∈* R^15599^) were reduced to 768 dimensions via principal component analysis to match the LitGene embedding (*E ∈* R^768^). These embeddings were then fused to create an integrated gene representation. Two approaches were employed to evaluate the predictive power of the fused embeddings in inferring gene-KEGG pathway relationships:

1. Pathway-Specific Logistic Regression: A separate logistic regression predictor was trained for each KEGG pathway. The fused features (*E ∈* R^1536^) were used as inputs to predict a binary label indicating the gene’s association with the pathway. This evaluation used a 5-fold cross-validation strategy, only considering pathways associated with at least 10 genes. For unbiased performance estimates, positive and negative gene-KEGG relationships were balanced for each KEGG pathway by randomly sampling an equal number of negative genes (i.e. genes not in the KEGG pathway).
2. Unified Logistic Regression: A single logistic regression classifier was used for all KEGG pathways. KEGG pathway embeddings from LitGene were concatenated with the fused gene embeddings (*E ∈* R^2304^) to predict gene-pathway associations. The same balancing strategy ensured an unbiased evaluation.

### Explaining Word Importance To LitGene Predictions

To elucidate the importance of specific words in LitGene predictions, we employed SHAP (SHapley Additive ExPlanations) values. SHAP values quantify the contribution of each input feature (e.g., sub-word tokens) to the model output relative to a baseline model output (e.g., all mask tokens) [52]. The calculation of SHAP values for NLP models is computationally expensive, so the SHAP library partition function was used to calculate Owen values [44], which considers that adjacent tokens (or players in the game-theoretic perspective) may form strategic coalitions. The partition function groups adjacent tokens and calculates their contribution as a whole, then attributes the coalition’s contribution uniformly across its members. Consequently, unimportant words may be grouped with important words, thus boosting their contribution or vice versa. The coalition size was dynamically adjusted using the hyperparameter for maximum coalition size.

SHAP values were calculated for various tasks, and the distribution of these values was analyzed across all samples to determine if certain words consistently held importance for specific tasks. As BERT may utilize sub-word tokens that are not always interpretable, the additive nature of SHAP values was leveraged; the sum of SHAP values for all features in a sample equals the difference between the baseline output and the actual model output. Consequently, we aggregated the contributions of all sub-word tokens within each word to obtain its total contribution.

Words were extracted from samples as consecutive characters in a sentence separated by spaces, including any attached punctuation. Distributions of words differing only by punctuation were combined. Additionally, distributions of words that were the plural forms of others in the set were merged. A wide distribution of SHAP values was observed for each word, primarily due to contextual variations. To account for this, the 90th percentile SHAP value was used as a characteristic measure of a word’s importance, capturing its relative importance.

To study LitGene’s capacity to understand and utilize biological text to predict gene properties, we employed the Cloze test, in which words are masked, and the model is tasked with predicting the masked words. The pre-trained weights of our model were loaded into a masked language model class, and out-of-distribution sentences were tested to fill the masked words. As a specific use case, we focused on the sentence: “Membrane proteins are more likely insoluble because their MASK residue interacts with the MASK layer of membrane.” This example allowed us to assess LitGene’s ability to comprehend and predict contextually appropriate words within a biomedical context.

### Baselines

The baselines that were used to benchmark LitGene in Table 1 consisted of representation-learning methods based on gene co-expression, single-cell transcriptome, or text. Below is a description of the baselines:

- **Majority Classifier**: The most frequent class in the dataset was predicted for all genes.
- **Gene2Vec** [17]: The Gene2Vec embeddings based on gene co-expression datasets.
- **Doc2Vec** [46] is a text-based embedding model that utilizes fixed-length feature representation to generate embeddings for variable-length text such as sentences, paragraphs, and documents. We get the embeddings by passing the gene summaries where we used an embedding size of 50, the maximum distance between the current and predicted word within a sentence of 2, all words with a total frequency of 1, and 40 training epochs.
- **LitGene-Base**: We also tested the effect of no fine-tuning on our model by generating the gene summary embeddings from BERT directly. We use BERT-PubMed base as our encoder, a 12-layer transformer-based model trained on PubMed data. We used CLS pooling and an embedding size of 768 to get the gene embeddings. We used Logistic regression to make the task predictions.
- **XLNet** [93] is an autoregressive pretraining transformer-based model. We use the 12-layer “xlnet-base-cased” and CLS pooling to get the gene embeddings.
- **PMC-LLaMA** [89] is a LLaMA-based [79] foundation language model that is pre-trained on the biomedical text and calibrated for medical domain applications. We get the embeddings from PMC-LLaMA by using Prompt-based last token pooling [37] where we use the following prompt *”This sentence: {text} means in one word:[CLS]”* and utilize the contextualized embedding of the last token which would be the CLS token that is added by the tokenizer after the colon.
- **GPT2** [65] is another open-source foundation language model that is trained on large text data and calibrated for downstream applications. We obtained the gene embeddings from the encoder of GPT2 by performing CLS pooling on the gene summaries.
- **Count Vectorizer** [59] which utilizes word counts as features for every gene in the dataset.
- **Term Frequency-Inverse Document Frequency (TF-IDF)** [88] which also utilizes word counts while considering their relative importance to the entire dataset.

The proposed baselines generate summary embeddings that we fine-tune for down-stream tasks by linear/logistic regression on top of the embeddings.

### Tasks and Datasets

To create the data displayed in Table 1, we benchmarked LitGene performance using the following tasks:

1. **Cytosolic Solubility.** Distinguishing between membrane proteins and cytosolic (soluble) proteins is important in proteomics. Soluble proteins are part of the cytosol, capable of traveling across membranes, and serve a wide range of functions both inside and outside of cells [19]. In contrast, membrane proteins play structural and functional roles in the cell as transporters, receptors, and enzymes. Over 50% of medications available on the market target membrane proteins, many of which have significant pharmacological implications [8]. We used LitGene to classify these protein types based on the textual description of their functions. To compile the dataset of soluble and insoluble genes, annotations were collected from two sources, resulting in a total of 988 soluble and 966 insoluble genes. The first data source utilized centrifugation assays to analyze protein solubility by measuring the extent of protein aggregation upon attempting dissolution [19]. The second data source employed in situ localization staining, classifying proteins as soluble if they were found in the lumen of organelles[2]. These two datasets were combined for the diversity of training and testing for the solubility prediction task. Given that centrifugation assays are more accurate for determining solubility, conflicts between the two sources were resolved in favor of the centrifugation data. Thus, the true positive set for soluble proteins included:

- All proteins identified as soluble in the centrifugation assay.
- Proteins classified as soluble in the location-based assay, excluding those labeled as insoluble in the centrifugation assay. Similarly, the true positive set for insoluble proteins included:
- All proteins identified as insoluble in the centrifugation assay.
- Proteins classified as insoluble in the location-based assay, excluding those labeled as soluble in the centrifugation assay.
2. **Chromatin State Prediction.** In epigenetic studies, chromatin states are identified by patterns of histone modification and DNA methylation, which not only provide a basis for genes to be segmented into biologically meaningful units but also help to identify the function of a given genomic segment in regulating gene expression. Bivalent Chromatin states, a hallmark of Embryonic Stem Cells (ESCs), are characterized by the presence of both repressive histone methylation (H3K27me3), and activating histone methylation (H3K4me3). This Bivalent Chromatin structure marks developmental genes and maintains their promoters in a poised state, ready for activation during differentiation processes [7]. We fine-tuned LitGene to classify genes with bivalent genomic regions, neither H3K4me3 nor H3K27me3 or were solely marked by H3K4me3, as reported by Theodoris et al. [76]. We utilized knowledge representations from gene summaries instead of single-cell transcriptomic data. We used 184 selected annotations available in the datasets for the bivalent and H3K4me3 classification task and 147 selected annotations available in the labeled datasets for the bivalent and unmethylated classification task from Theodoris et al. [76] al to fine-tune LitGene.
3. **Dosage Sensitivity.** Specific genes are said to be dosage sensitive when the variations in gene dosage (copy number variations) can cause phenotypic changes. For this task, we evaluate our model on a curated dataset of two annotations, namely dosage-sensitive and dosage-insensitive genes, from previously reported studies [76, 48, 71, 56].
4. **Subcellular Localization.** We fine-tuned LitGene to distinguish between the subcellular localization of each protein product of genes. We utilize a set of gene annotations that spans the following 3 annotations: Cytoplasm, Cell membrane, and Nucleus, from the UniProt database, as described in Almagro et al. [2]. Understanding protein subcellular locations is essential for understanding its function and physiochemical properties. It can help identify possible targets for therapy and understand illnesses associated with abnormal subcellular localization [77].
5. **Conservation.** The PhastCon score is a measure derived from the PHAST (Phylogenetic Analysis with Space/Time models) package. It quantifies the evolutionary conservation of genomic sequences, offering insights into the functional significance and evolutionary pressures shaping these regions. Predicting PhastCon scores is essential for identifying functionally important genomic elements, such as coding sequences and regulatory regions, which are critical in evolutionary biology, functional genomics, and disease studies [66]. LitGene performance to predict Gene Conservation is measured by predicting the PhastonCon score with the help of knowledge representations of gene summaries obtained from LitGene as input. The Spearman and Pearson correlation coefficients are used to evaluate this regression task.
6. **Transcription Factors (TFs) Range.** TFs are the proteins that bind to certain sequences of DNA and control the transcription of genetic information from DNA to messenger RNA [66], with their influence ranging from short-range effects on nearby genes to long-range effects influencing distant genes. The dataset spans two different annotations, namely short-range and long-range genes, from Theodoris et al. [76].
7. **Transcription Factors (TFs) Target Type.** We also study two specific transcription factors, GATA4 and TBX5 since they have crucial roles including heart development and function [76].We utilize annotations of genes that identify whether a gene is directly/indirectly regulated by one or both of these transcription factors. More specifically, we collect dataset annotations that span the following 5 labels: gata4_indirect, gata4_direct, tbx5_indirect, tbx5_direct, and combo_targets, Theodoris et al. [76].

## Data Availability

The data used for training including gene summaries, GO term summaries, gene-GO annotations, and task-specific annotations are freely available from public sources as listed in the Methods section.

## Code Availability

The data preprocessing scripts, source code for training, pre-trained model, tokenizer, and interpretability are available at https://github.com/vinash85/LitGene.git. We also release a web interface of LitGene at litgene.tumorai.org.

## A Supplementary Figures

**Supplementary Figure 1:**
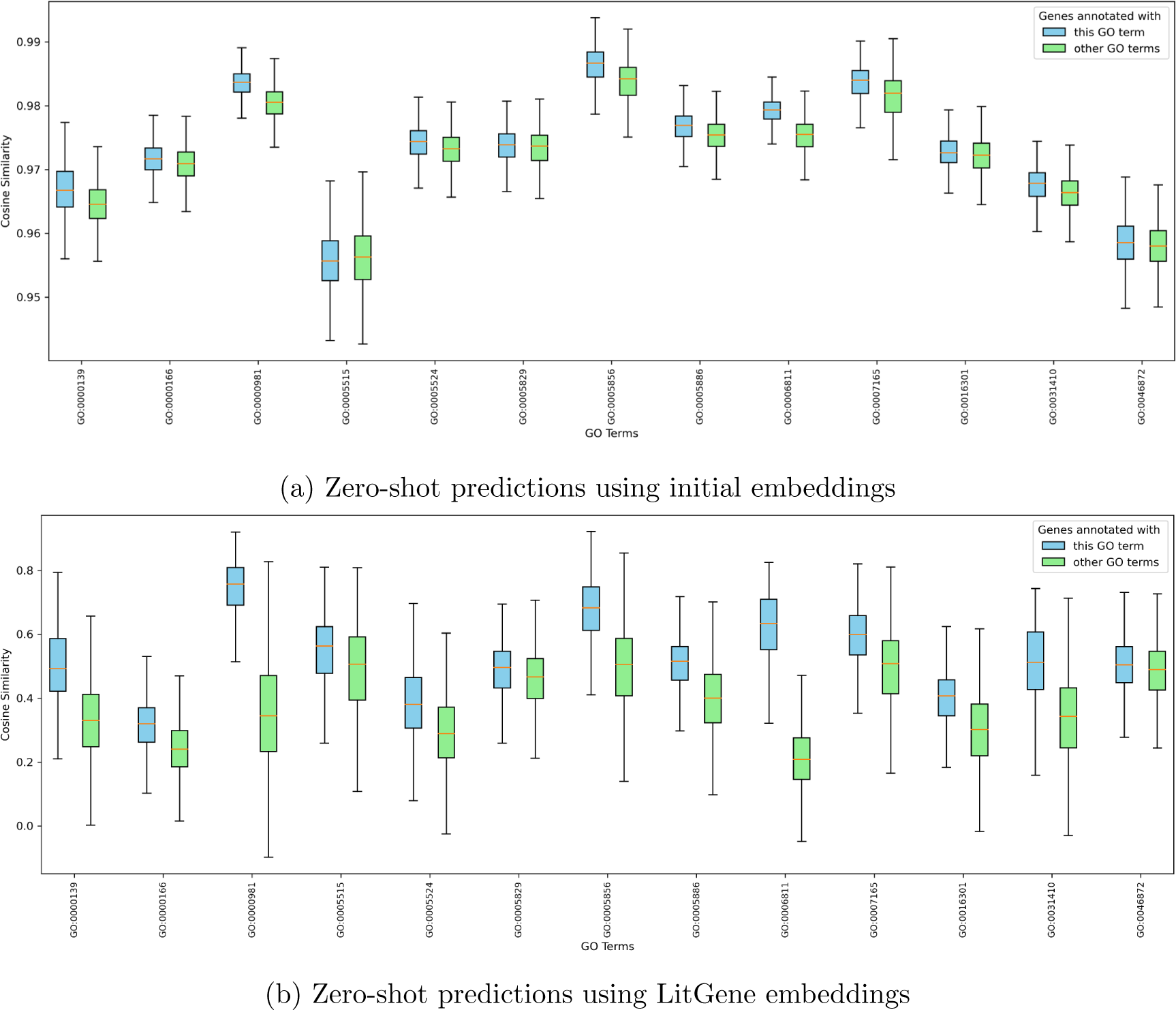
Expanded plots of the discovered novel gene-to-GO term relationships by employing contrastive learning (CL). Be aware that the scales of the two figures are not the same. LitGene-Base (i.e. initial embeddings of LitGene before applying CL; a) restricts the similarities within a narrower range, making the results less comparable, whereas LitGene-CL (i.e. LitGene after applying CL; b) spreads the gene similarities throughout the entire y-axis, enhancing the distinction between GO terms and genes.

**Supplementary Figure 2:**
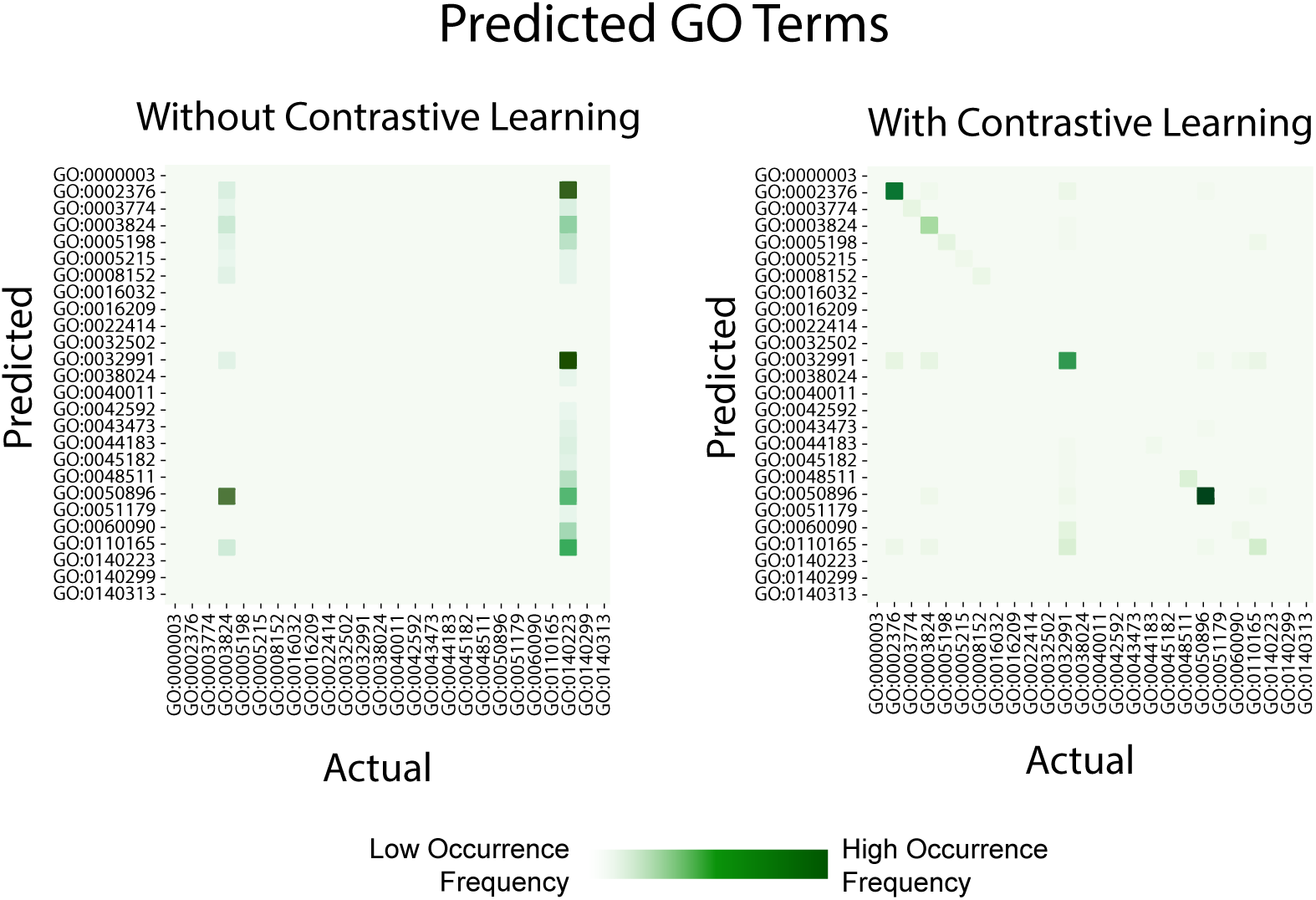
Confusion matrix of predicted GO relationships derived via naive thresholds on the similarities found in Supplementary Figure 1. (Left) The predictions of LitGene-Base: without utilizing contrastive learning and (Right) The predictions of LitGene-CL: after applying contrastive learning. In both plots, the x-axis/y-axis is the actual/predicted GO relationships for the genes belonging to that GO.

**Supplementary Figure 3:**
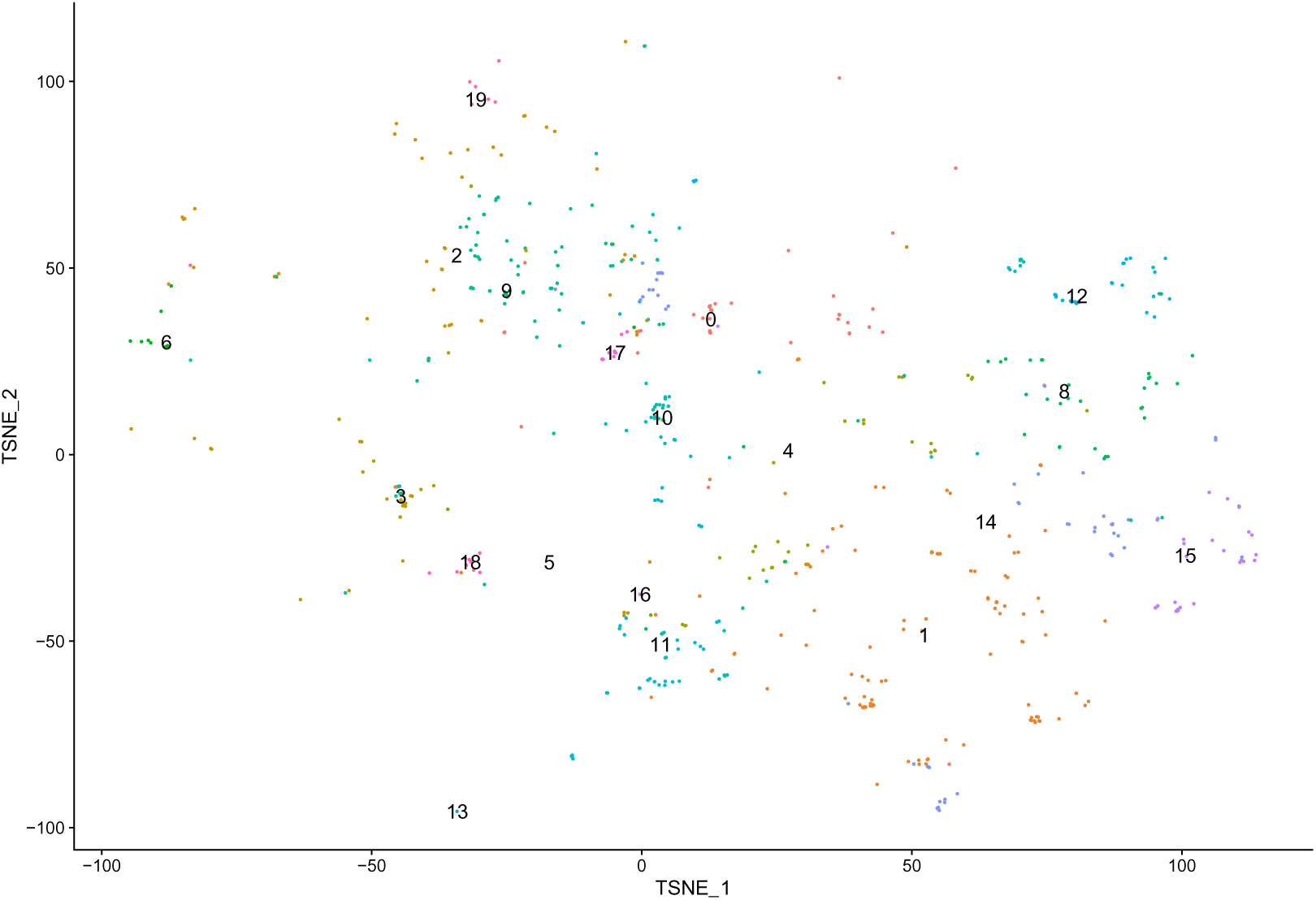
TSNE clustering of the reduced disease embeddings.

**Supplementary Figure 4:**
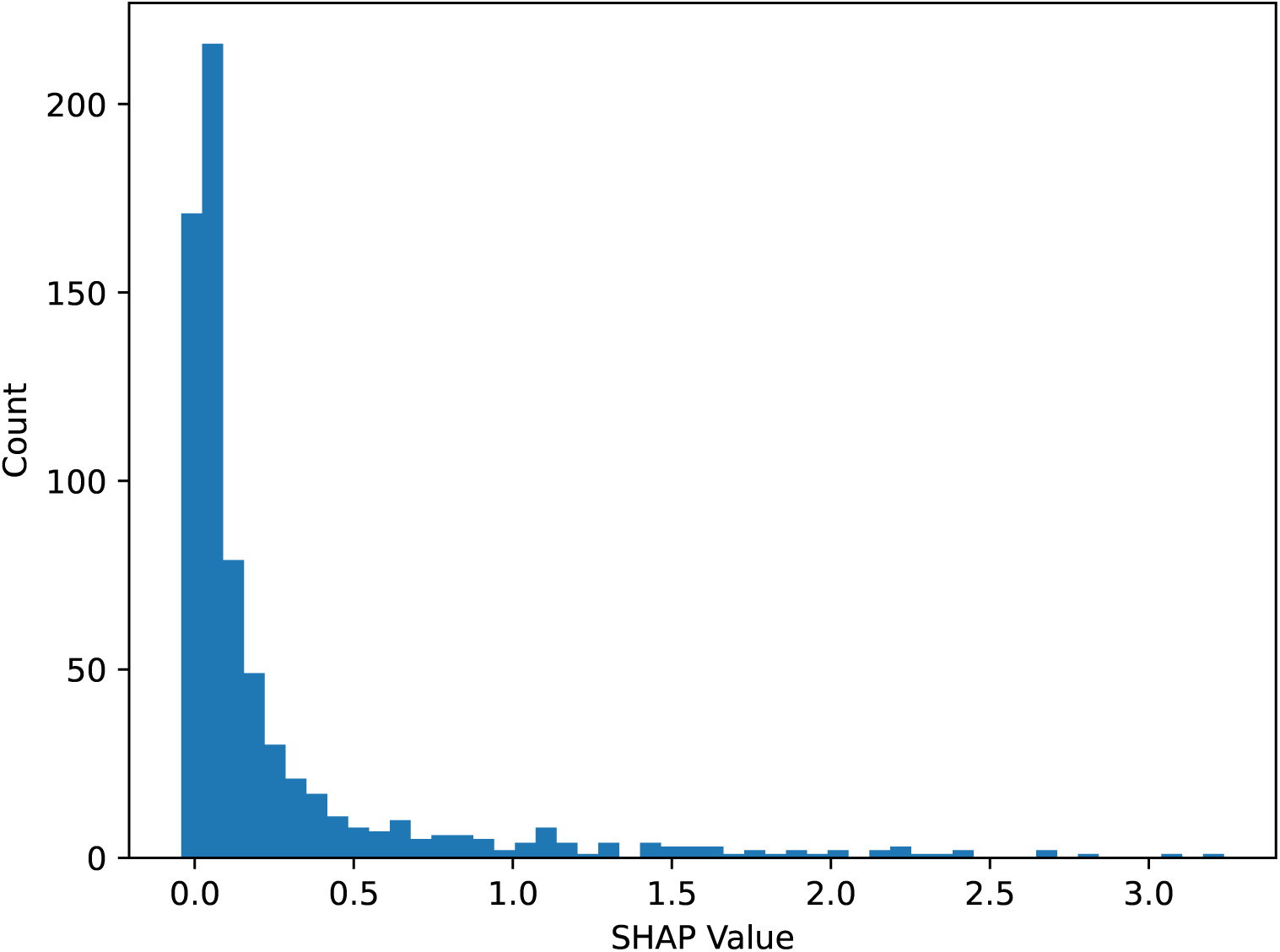
Histogram of SHAP values for the word *membrane* in the solubility training set. The training set contains 1499 gene summaries, in which 845 instances of the word *membrane* exist across 478 gene summaries.

**Supplementary Figure 5:**
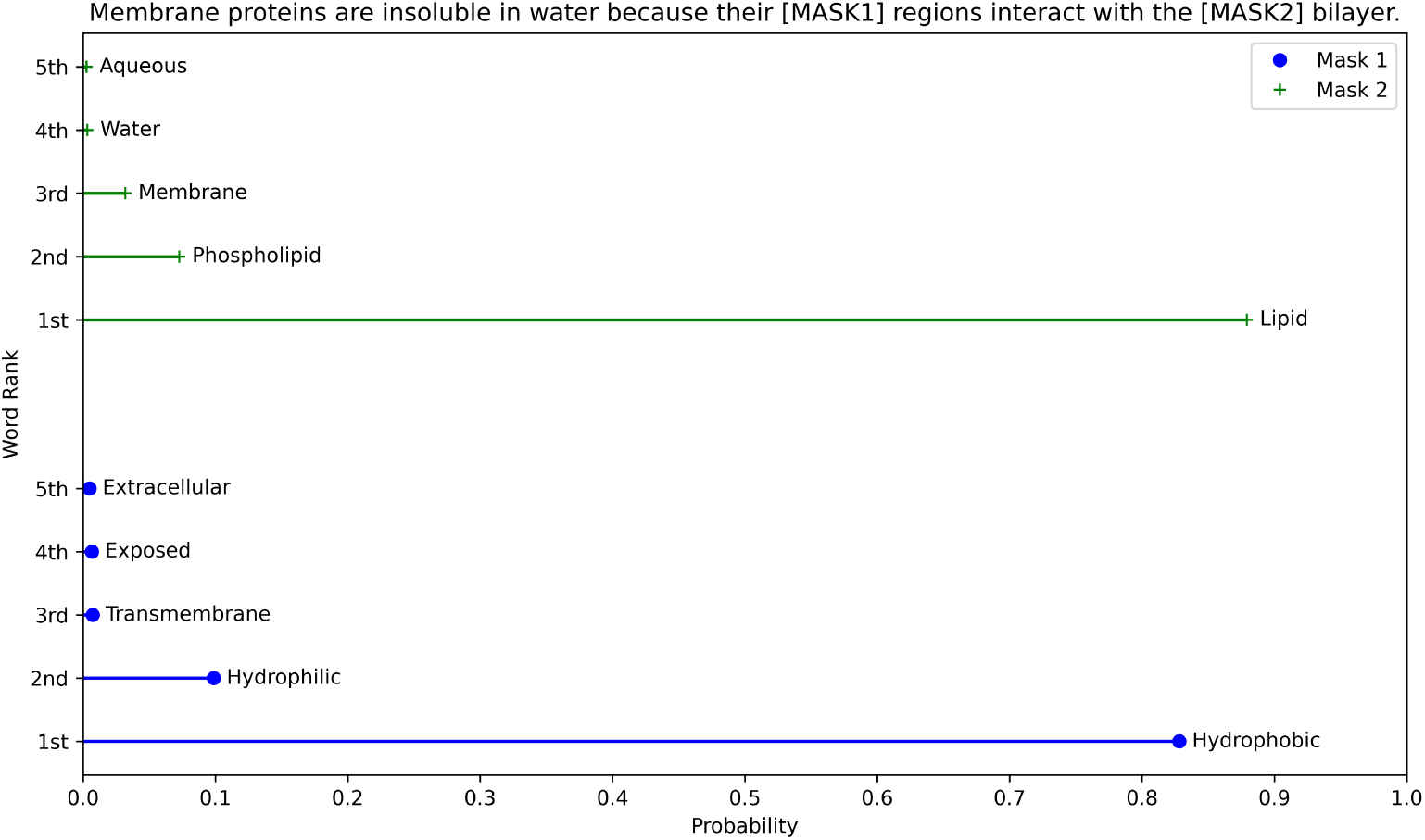
Cloze test performed to measure LitGene’s ability to internalize biological concepts. The sentence consists of two masked tokens where LitGene is asked to predict these tokens. The x-axis demonstrates the probability of the top 5 predicted tokens for each mask while the y-axis ranks the predicted tokens based on the probability.

**Supplementary Figure 6:**
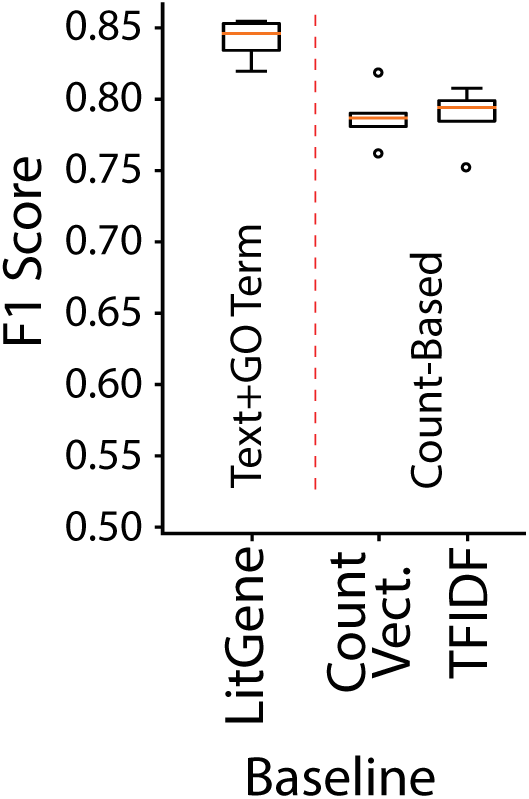
Evaluating data leakage of LitGene by comparing the performance to count-based feature extraction models, Count Vectorizer, and TF-IDF. The figure shows the performance of fine-tuning LitGene on the solubility task.

**Supplementary Figure 7:**
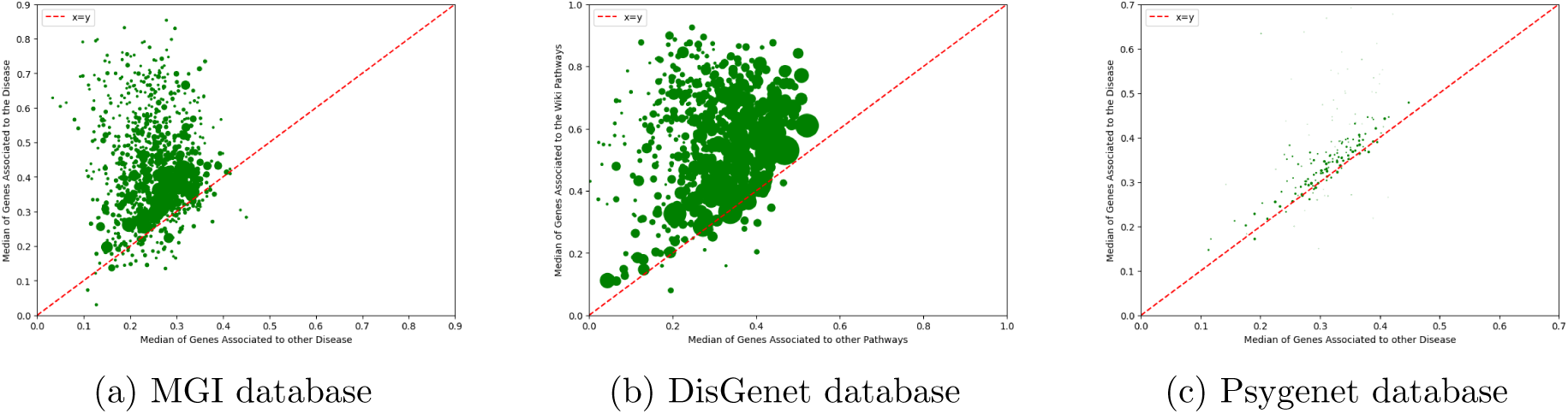
Gene-disease association results on other databases. The figure features three databases other than those mentioned in the main text. The scatter plot compares the median cosine similarity of genes associated with a specific disease to those associated with other diseases across different databases. Points above the x = y line suggest a stronger association with the specific disease. Note that the number of disease associations found in every database varies significantly.

**Supplementary Figure 8:**
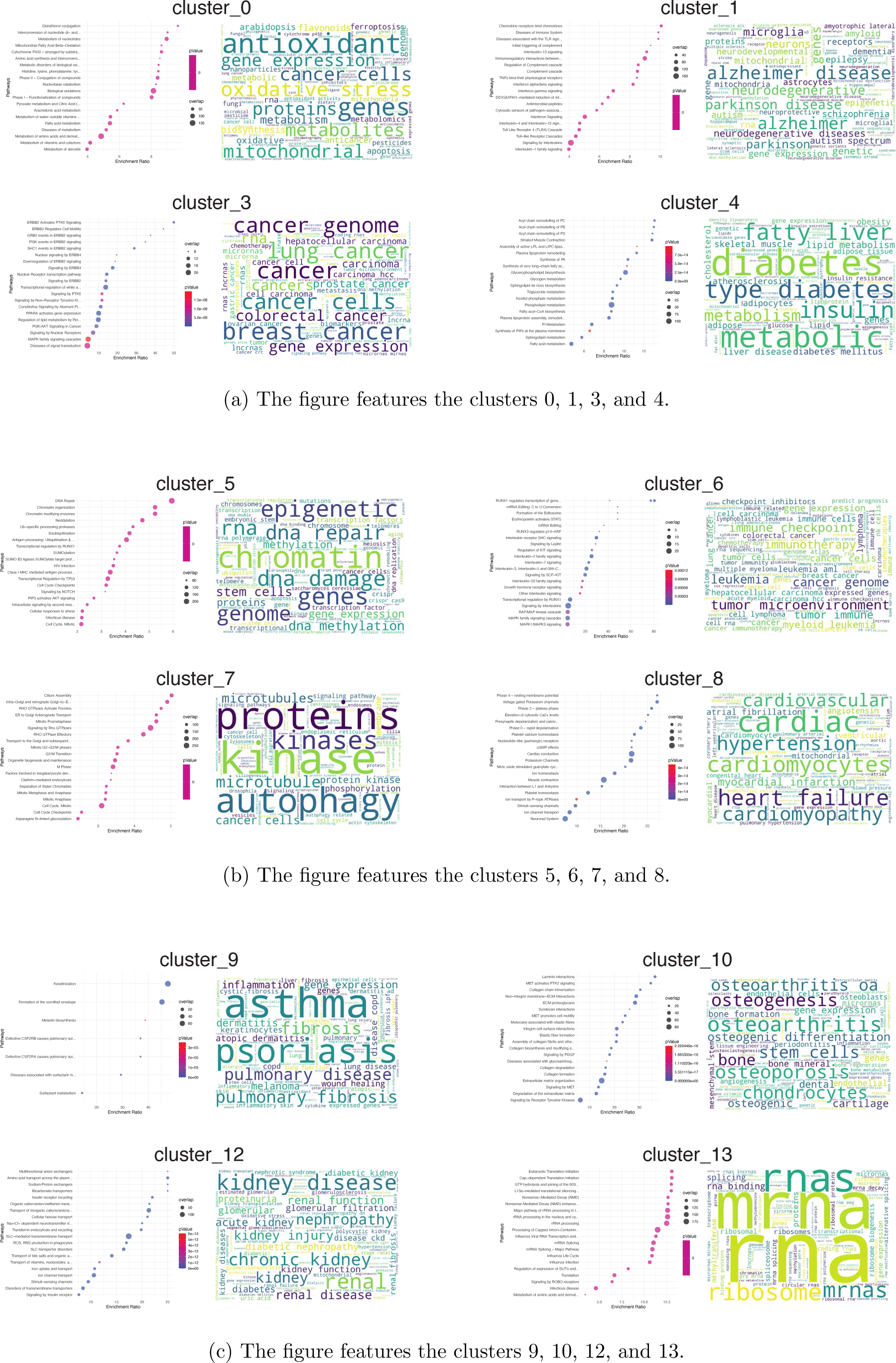
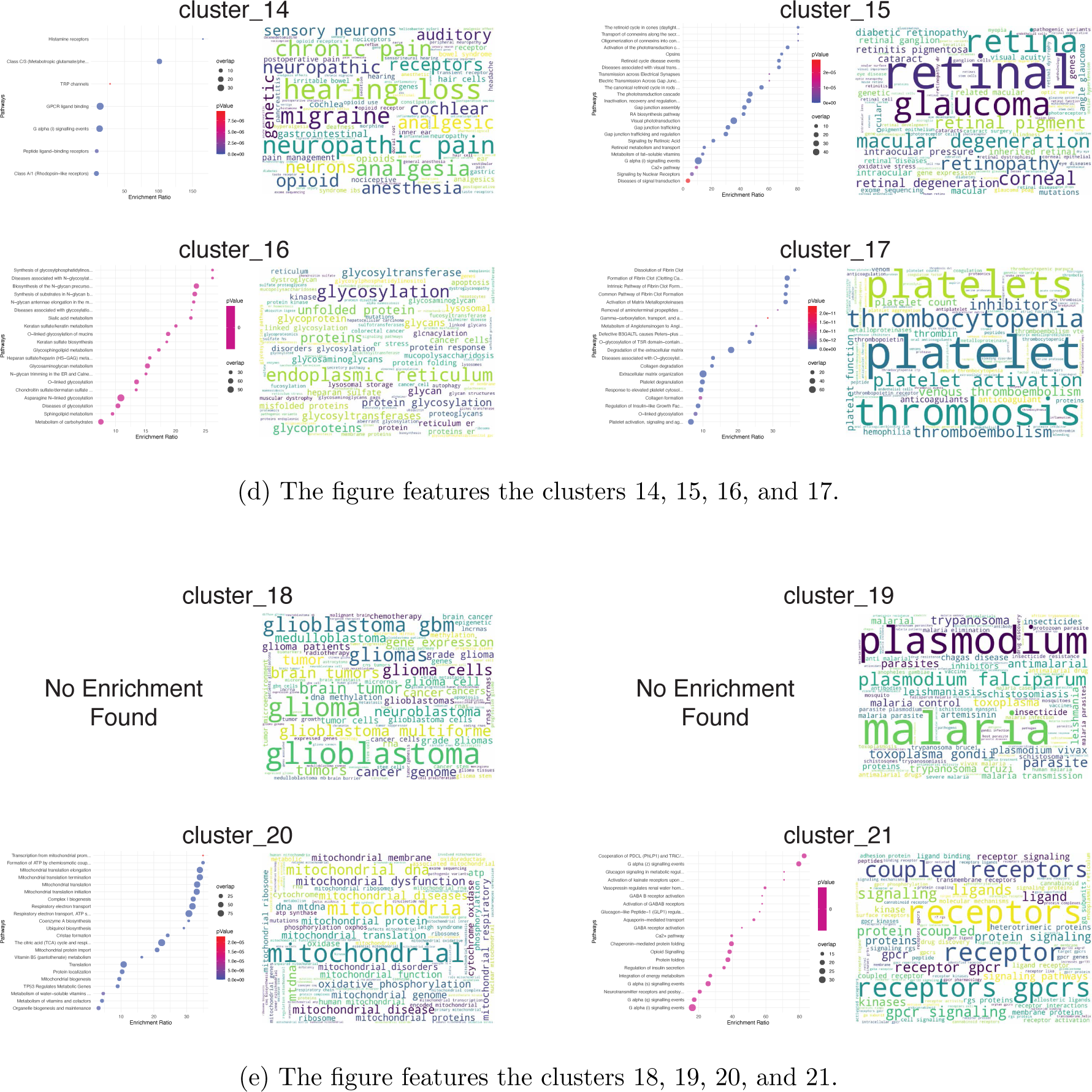
Anchoring Gene-Disease Predictions in Published Research. Various clusters are visualized across multiple figures to show the enrichment ratio and word clouds of topics extracted from articles.

## B Supplementary Tables

**Supplementary Table 1:** 5-fold Accuracy outcomes for contrastive learning-enhanced gene embeddings across a spectrum of gene prediction tasks.

| Model | Dosage<br>Sensitivity | BivalentVs<br>Lys4<br>Methylated | BivalentVs<br>Non<br>Methylated | Tf<br>range | Tf<br>target type | Solubility | Subcellular<br>localization | Conservation<br>(Pearson Corr.) |
| --- | --- | --- | --- | --- | --- | --- | --- | --- |
| Majority Classifier | 0.73 ± — | 0.58 ± — | 0.75 ± — | 0.73 ± — | 0.41 ± — | 0.52 ± — | 0.39 ± — | — ± — |
| GPT2 | 0.74 ± 0.04 | **0.86 ± 0.04** | 0.80 ± 0.11 | 0.71 ± 0.03 | 0.18 ± 0.02 | 0.80 ± 0.02 | 0.77 ± 0.01 | 0.31 ± 0.02 |
| Doc2Vec | 0.74 ± 0.04 | 0.84 ± 0.06 | 0.78 ± 0.05 | 0.66 ± 0.07 | 0.26 ± 0.01 | 0.71 ± 0.03 | 0.69 ± 0.02 | 0.34 ± 0.01 |
| PMC-LLaMA | 0.86 ± 0.05 | 0.77 ± 0.04 | **0.84 ± 0.07** | 0.64 ± 0.08 | 0.08 ± 0.01 | 0.78 ± 0.03 | 0.69 ± 0.01 | **0.55 ± 0.01** |
| XLNet | 0.74 ± 0.06 | 0.84 ± 0.06 | 0.83 ± 0.08 | 0.69 ± 0.05 | 0.12 ± 0.01 | 0.79 ± 0.02 | 0.76 ± 0.01 | 0.40 ± 0.01 |
| Gene2Vec | 0.84 ± 0.04 | 0.84 ± 0.06 | 0.75 ± 0.06 | **0.75 ± 0.08** | 0.21 ± 0.01 | 0.56 ± 0.02 | 0.54 ± 0.02 | 0.50 ± 0.02 |
| CountVectorizer | 0.79 ± 0.03 | 0.84 ± 0.02 | 0.76 ± 0.05 | 0.65 ± 0.05 | 0.07 ± 0.04 | 0.68 ± 0.20 | 0.78 ± 0.01 | 0.05 ± 0.015 |
| TF-IDF | 0.80 ± 0.04 | 0.87 ± 0.04 | 0.61 ± 0.01 | 0.62 ± 0.02 | 0.07 ± 0.03 | 0.68 ± 0.19 | 0.80 ± 0.01 | 0.03 ± 0.02 |
| LitGene - no Finetuning | 0.76 ± 0.09 | 0.83 ± 0.06 | 0.77 ± 0.10 | 0.68 ± 0.04 | 0.17 ± 0.01 | 0.76 ± 0.02 | 0.76 ± 0.01 | 0.48 ± 0.02 |
| LitGene | **0.87 ± 0.06** | **0.86 ± 0.09** | 0.82 ± 0.08 | 0.74 ± 0.07 | **0.49 ± 0.04** | **0.89 ± 0.01** | **0.83 ± 0.01** | 0.53 ± 0.01 |

**Supplementary Table 2:** A comprehensive evaluation of gene products solubility of solubility specific models from DeepSoluE (Wang et al., 2023).

| Model | F1 Score | Accuracy |
| --- | --- | --- |
| RPSP | 0.392 | 0.498 |
| ccSOL omics | 0.537 | 0.508 |
| SKADE | 0.168 | 0.492 |
| SOLpro | 0.468 | 0.52 |
| Protein-Sol | 0.585 | 0.516 |
| DeepSol | 0.239 | 0.529 |
| rWH | 0.485 | 0.54 |
| ESPRESSO | 0.583 | 0.538 |
| CamSol | 0.487 | 0.541 |
| SWI | 0.638 | 0.559 |
| PROSO II | 0.491 | 0.58 |
| SoluProt | 0.593 | 0.585 |
| DeepSoluE | 0.600 | 0.5952 |
| LitGene - no Finetuning | 0.770 | 0.763 |
| LitGene | **0.843** | **0.832** |

**Supplementary Table 3:** Confusion matrix for gene summaries in the test set of the annotated solubility dataset, where the i-th row represents the true class and the j-th column represents the predicted class.

| Containing 'membrane' | Insoluble | Soluble |
| --- | --- | --- |
| Insoluble | 96 | 18 |
| Soluble | 8 | 28 |

| Without 'membrane' | Insoluble | Soluble |
| --- | --- | --- |
| Insoluble | 84 | 42 |
| Soluble | 23 | 190 |

## C Website (litgene.tumorai.org)

**Supplementary Figure 9:**
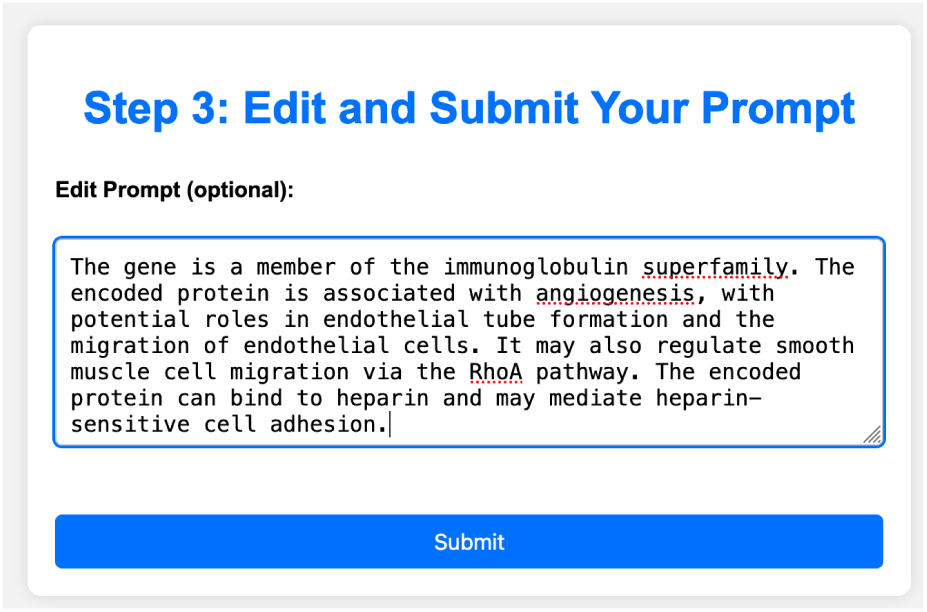
Input prompt. After selecting an input mode (gene/disease/drug) and a specific instance, the website generates a default text input for analysis. Shown here is the default description of the AAMP gene, which can optionally be edited by the user.

**Supplementary Figure 10:**
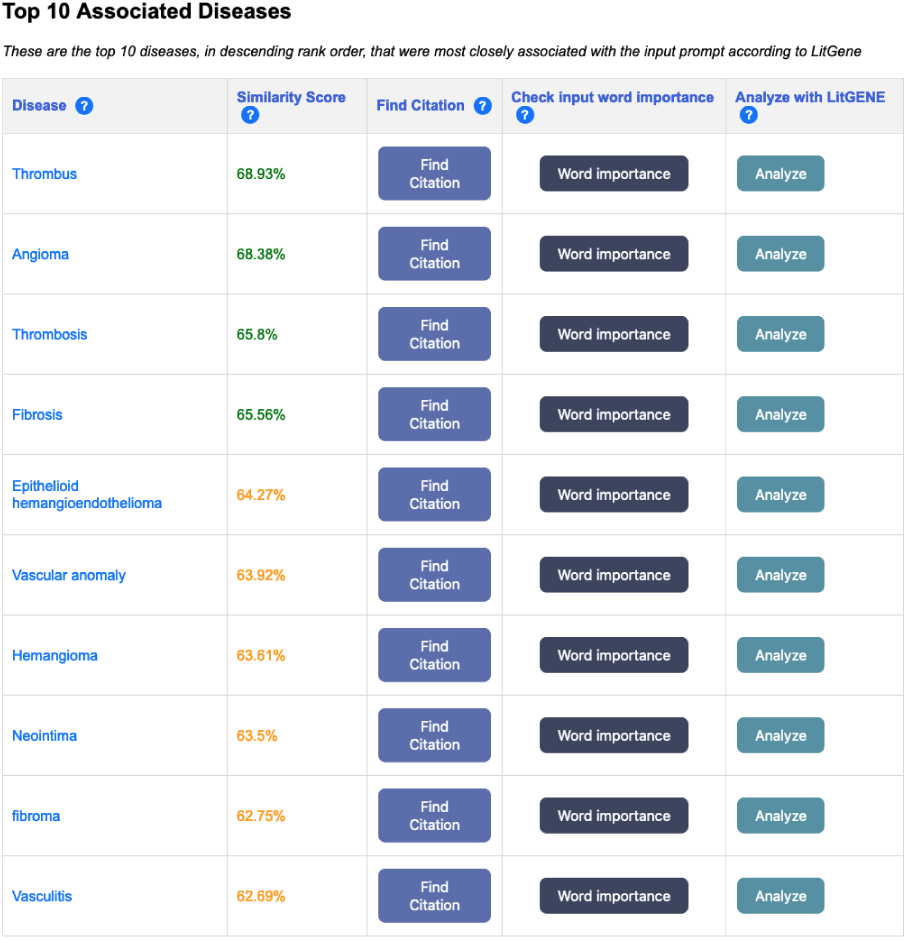
Output. After submitting an input to the website, the top 10 associated items of each mode are listed (gene/disease/drug). Shown above are the top 10 diseases associated to the AAMP gene input.

**Supplementary Figure 11:**
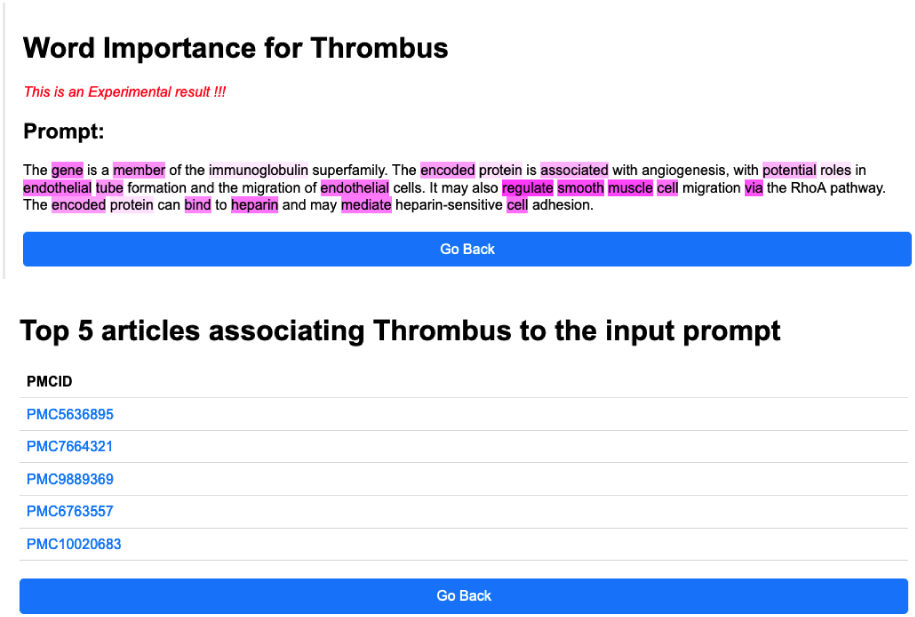
Analyses. (Top) The word importance analysis for disease association to the AAMP gene. Words are highlighted according to their SHAP value, measuring each word’s contribution to the cosine similarity between the input prompt and the default text description of Thrombus. (Bottom) The top 5 articles most related to Thrombus and the input gene description of AAMP as measured by cosine similarity to the corpus of PubMed abstracts.

